# Enhanced splicing modulation by NMA-modified antisense oligonucleotides

**DOI:** 10.1101/2025.10.06.680653

**Authors:** Karen Ling, Thazha P. Prakash, Jinghua Yu, Michaela Jackson, Seung J. Chun, Gemma Bachmann, Noah Post, Sarah Greenlee, Armand Soriano, John L. Hunyara, Daniel A. Norris, Paymaan Jafar-nejad, Eric E. Swayze, C. Frank Bennett, Frank Rigo

## Abstract

Aberrant RNA splicing contributes to many human diseases, and splice-switching antisense oligonucleotides (SSOs) are ideally suited as a therapeutic strategy to modulate splicing and restore normal gene expression. Nusinersen (Spinraza™) has revolutionized the treatment of spinal muscular atrophy. It is a splice-switching oligonucleotide (SSO) that is modified with 2’-*O*-methoxyethyl (MOE) modifications. Here, we introduce a next-generation ribose modification, 2′-*O*-[2-(methyl-amino)-2-oxoethyl] (NMA), which enhances the pharmacological properties of SSOs. We identified a long-lasting NMA-modified human candidate SSO, salanersen, that is 3-4-fold more potent than nusinersen in human *SMN2* transgenic mice. To evaluate the generality of the NMA chemistry, we applied it to modulation of *SCN1A* exon 20N splicing, a therapeutic strategy for Dravet syndrome. An NMA-modified SSO is 3.5-fold more potent than STK-001, a MOE-modified SSO currently in clinical trials. Our data establish the NMA chemistry as a broadly applicable ribose modification that markedly improves the pharmacological profile of SSOs, supporting its development as a next-generation platform for splicing modulation therapies.

**Graphical abstract:** 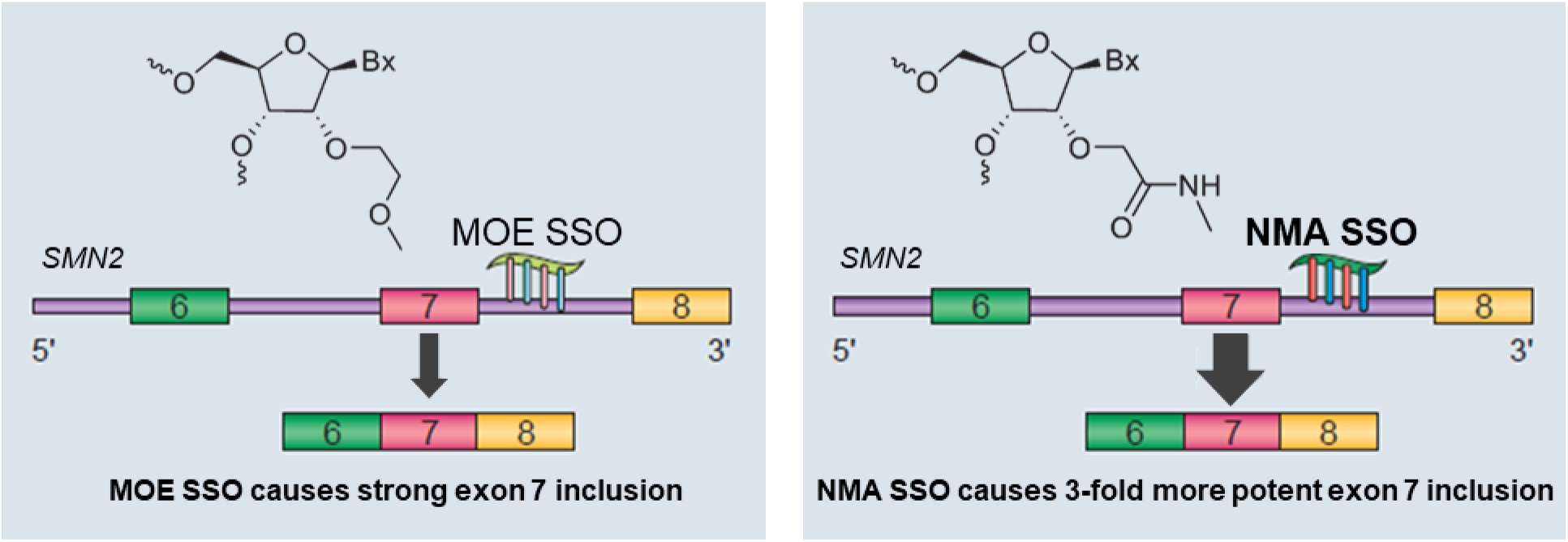

## Introduction

Alternative splicing is ubiquitous in eukaryotic cells and expands the proteomic output of the genome^1,2^. Mutations in cis-acting splicing elements or in trans-acting splicing factors that disrupt splicing and frequently contribute to many human diseases. Splice-switching oligonucleotides (SSOs) have emerged as a therapeutic strategy to correct splicing defects or modulate gene expression by redirecting splicing^3^. SSOs are short (15–25 nt) antisense oligonucleotides that are chemically modified throughout their length. They hybridize to pre-mRNA and redirect splicing by sterically blocking splicing factors. SSOs have been used successfully to modulate splicing for many targets in a variety of cell types and tissues and many SSOs have advanced into individualized N-of-1 studies as well as broader clinical trials^4–9^ . Furthermore, several SSOs have received marketing authorization^10–15^ . In the central nervous system (CNS), two diseases where SSOs have advanced the furthest are spinal muscular atrophy (SMA) and Dravet syndrome.

SMA is a hereditary neurodegenerative disease characterized by the loss of motor neurons, progressive muscle weakness, and atrophy^16^. This condition is caused by homozygous loss-of-function mutations in the SMN1 gene, leading to a deficiency of the SMN protein^17,18^. Although the paralogous gene *SMN2* also produces the SMN protein, its transcripts yield only 5–10% of the functional, full-length protein. The limited production of functional protein from *SMN2* is largely due to a C-to-T transition that disrupts splicing regulation, promotes exon 7 skipping, and results in a truncated, nonfunctional protein^19–21^. Nusinersen (Spinraza™) is a 2′-*O*-methoxyethyl (MOE) modified SSO that binds ISS-N1, an intronic splicing silencer in intron 7 of *SMN2* pre-mRNA^22^. By masking ISS-N1, nusinersen prevents the binding of splicing repressors (notably hnRNP A1/A2), thereby promoting exon 7 inclusion and yielding full-length SMN protein^23^. Nusinersen delivered strongly positive results in two randomized, sham-controlled trials for patients with spinal muscular atrophy^10,11^. In infants with devastating early-onset disease, treatment markedly increased survival without permanent ventilation and substantially improved motor milestone attainment^10^ . In children with later-onset SMA, nusinersen led to robust improvements in overall motor function, while untreated patients experienced progressive decline^11^.

The success of nusinersen has propelled efforts to test SSOs in additional CNS diseases, including Dravet syndrome which is a severe developmental and epileptic encephalopathy most often caused by heterozygous loss-of-function mutations in the *SCN1A* gene, resulting in haploinsufficiency of the voltage-gated sodium channel Nav1.1. The *SCN1A* gene produces an abundant splice isoform that includes a poison exon known as exon 20N. Inclusion of exon 20N introduces a premature termination codon (PTC), leading to transcript degradation through nonsense-mediated decay (NMD)^24,25^. Modulating splicing to reduce inclusion of exon 20N in transcripts from the functional *SCN1A* allele prevents NMD, thereby increasing the levels of *SCN1A* mRNA and Nav1.1 protein. Preclinical studies in a mouse model of Dravet syndrome have demonstrated that SSOs designed to exclude exon 20N increase the levels of Nav1.1 protein, reduce seizures, and extend survival^26,27^. Based on these findings, a MOE-modified SSO (STK-001), has advanced into clinical evaluation in Dravet syndrome patients, with promising early results (ClinicalTrials.gov ID: NCT04740476)^28^.

In mouse models of SMA, several types of SSOs have been tested, including those based on 2′-*O*-methyl (2′-OMe), phosphorodiamidate morpholino oligomers (PMOs), and 2′-*O*-methoxyethyl (MOE) chemistries. When administered to the CNS, SSOs with these chemistries promote exon 7 inclusion, increase SMN protein expression, and extend survival^29–33^. In comparative dose– response studies in *SMN2* transgenic mice, MOE SSOs were more potent at promoting exon 7 inclusion than PMO or 2′-OMe SSOs. Constrained ethyl (cEt)–modified oligonucleotides, evaluated in *SMN2* transgenic mice, showed activity but no clear advantage over MOE^34^. By contrast, an SSO with 2′-fluoro modifications unexpectedly induced *SMN2* exon 7 skipping by recruiting the RNA-binding proteins ILF2/ILF3^35^. Additional chemistries, such as tricyclo-DNA (tcDNA) have also shown activity in mice^36^, but MOE remains the chemistry with the most advanced clinical progress for splicing modulation. Thus, it is the only chemistry that has been explored for modulating *SCN1A* exon 20N splicing for the treatment of Dravet syndrome ^26^ .

Despite the success of 2’-MOE modified SSOs for CNS disease, advances in oligonucleotide chemistries provide an opportunity to further enhance the profile of SSOs. More potent SSOs should result in better clinical response at lower doses, potentially improving their safety profile. In addition, decreasing the dosing interval would be highly desirable for intrathecal administered SSOs. To address this need we investigated novel chemical modifications not previously applied to SSOs, specifically 2ʹ-*O*-[2-(methyl-amino)-2-oxoethyl] (NMA) and its analogs. Here we describe the synthesis, optimization, and pharmacological characterization of NMA modified SSOs targeting *SMN2* and *SCN1A*.

## Results

### Enhanced potency of an NMA-modified SMN2 SSO

Pre-mRNA splicing of the *SMN2* gene predominantly generates a transcript lacking exon 7 and results in an unstable SMN protein that is rapidly degraded (Fig. 1A). Nusinersen, an SSO uniformly modified with MOE promotes exon 7 inclusion and increases SMN protein levels. We set out to identify a chemical modification with improved *in vivo* potency (i.e., more activity at a lower dose) for *SMN2* exon 7 inclusion compared to nusinersen. A series of 2’-*O*-[2-(amino)-2-oxoethyl]–modified SSOs have previously been shown to have high binding affinities to complementary RNA, comparable to the MOE modification^37^. Within this class of 2’-*O*-substituted sugar modifications, the *N*-methylacetamide (NMA) modification emerged as a distinct analog, displaying structural features reminiscent of MOE in crystallographic analyses, but with superior metabolic stability^37,38^. Increasing doses of the uniform MOE SSO nusinersen (here referred as MOE-1), a uniform NMA SSO (NMA-1), or SSOs modified with analogs of NMA (Fig. 1B; Supplementary information, Table S1) with varied steric and electronic characteristics, were administered to the cerebrospinal fluid (CSF) of adult human *SMN2* transgenic mice^39^ via a single intracerebroventricular (ICV) bolus injection. Brain and spinal cord tissues were analyzed for *SMN2* splicing 14 days post-injection by real-time RT-PCR. All SSOs induced dose-dependent *SMN2* splicing correction, evidenced by an increase in full-length (exon 7–included) transcripts and a corresponding decrease in Δ7 (exon 7–skipped) transcripts in both the spinal cord (Fig. 1C) and the brain (Extended Data Fig. 1). Based on the half-maximal effective dose (ED_50_) for promoting *SMN2* exon 7 inclusion, the NMA-modified SSO (NMA-1) emerged as the most potent, exhibiting the lowest ED_50_ value among all SSOs tested.

**Figure 1.**
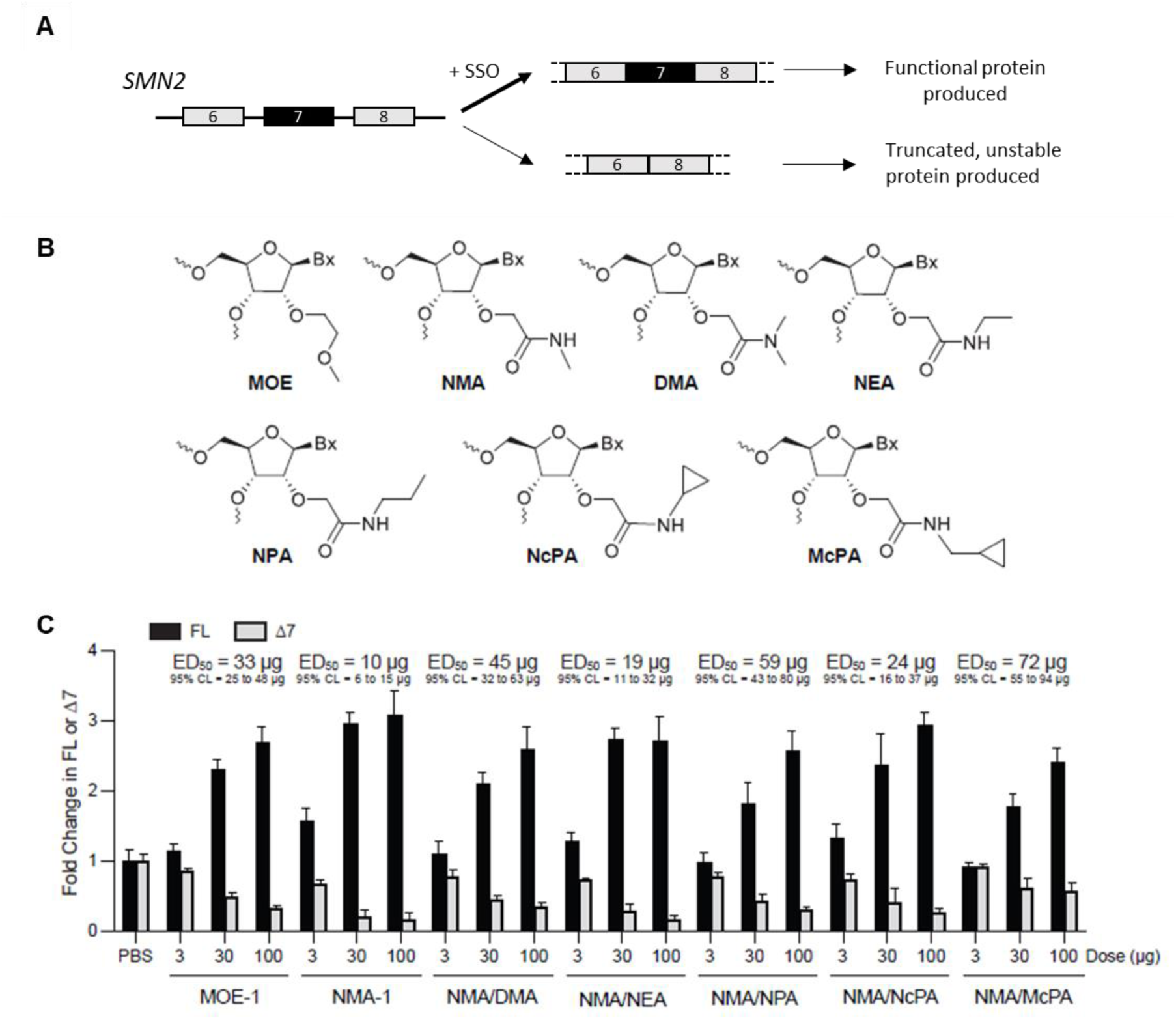
Potency for *SMN2* splicing correction of SSOs modified with MOE, NMA or NMA analogs in the spinal cord of the human *SMN2* transgenic mice. (A) Schematic diagram of naturally occurring productive vs nonproductive alternative splicing of *SMN2* pre-mRNA. The *SMN2* transcript lacking exon 7 encodes a truncated SMN protein that is unstable and rapidly degraded. Splice-switching oligonucleotide (SSO) promotes inclusion of exon 7 generating a full-length mRNA and functional protein. (B) Chemical structures of nucleotides modified with MOE, NMA, and NMA analogs. (C) Real-time RT-PCR analysis of *SMN2* transcripts including exon 7 (FL) or skipping exon 7 (Δ7) in the spinal cord 14 days after intracerebroventricular (ICV) bolus injection of MOE-1, NMA-1 or SSOs modified with analogs of NMA in *SMN2* transgenic mice. For each dose level, n=4. Error bars represent the S.D. The ED_50_ for exon 7 inclusion (FL) and confidence interval (CI) are shown.

To more precisely compare the potency for *SMN2* splicing correction of the NMA versus MOE modified SSOs, we performed a five-point dose-response of MOE-1 and NMA-1 for *SMN2* correction in *SMN2* transgenic mice. *SMN2* splicing correction was measured in the spinal cord and brain harvested 2 weeks after the ICV bolus injection by real-time RT-PCR. The half-maximal effective dose (ED_50_) for *SMN2* splicing correction by the NMA-1 SSO was 5 µg in the spinal cord and 9 µg in the brain, and the ED_50_ for *SMN2* splicing correction by the MOE-1 SSO was 22 µg in the spinal cord and 32 µg in the brain (Extended Data Fig. 2A,D), showing that NMA modification was 3- to 4-fold more potent than the MOE modification.

To determine if the increased potency of the NMA-1 SSO observed in *SMN2* transgenic mice was due to greater accumulation in CNS tissues, we measured the amount of SSO in CNS tissues. The tissue concentration of the NMA-1 SSO was not higher than that of the MOE-1 SSO in the spinal cord and brain (Extended Data, Fig. 2B,C,E,F). We also examined the relationship between the amount of the NMA-1 SSO in the CNS tissue and the degree of *SMN2* splicing correction. For each mouse that was dosed in Extended Data Fig. 2A&D, we measured SSO concentration in the spinal cord and brain, and plotted these values against the degree of *SMN2* splicing correction in the spinal cord and brain for the same mouse. We observed a strong correlation between SSO levels in CNS tissue and *SMN2* splicing correction. The half-maximal effective concentration (EC_50_) of the NMA-1 was ∼5 times lower than the MOE-1 SSO (0.3 µg/g vs. 1.5 µg/g in the spinal cord (Extended Data Fig. 2B,C) and 1.1 µg/g vs 5.3 µg/g in brain tissues (Extended Data Fig. 2E,F). Therefore, even though both the MOE-1 and NMA-1 SSOs achieved similar CNS tissue concentrations, the NMA-1 SSO was approximately 5-fold more active for *SMN2* splicing correction than the MOE-1 SSO, consistent with its higher potency on an ED_50_ basis.

Since the NMA-1 SSO showed enhanced potency in CNS tissues, we determined if this was also the case in peripheral tissues. Increasing doses were administered subcutaneously to adult *SMN2* transgenic mice on days 1, 3, 5 and 7. Several peripheral tissues were analyzed for *SMN2* splicing correction 3 days after the last dose by real-time RT-PCR. In all tissues examined, both the NMA-1 and the MOE-1 SSOs resulted in a dose-dependent increase in *SMN2* splicing correction. The NMA-1 SSO was >2.5-fold more potent than the MOE-1 SSO for *SMN2* splicing correction (Extended Data Fig. 3). The potency of the NMA-1 SSO for *SMN2* splicing correction in peripheral tissues was further improved by ∼2-3-fold when conjugated to palmitic acid (C16) (Extended Data Fig. 4). This suggests that alternative conjugation strategies for enhanced delivery to a variety of cell types and tissues^40^ should be additive to the NMA modification. Additionally, the potency enhancement conferred by the NMA modification is distinct from C16-mediated protein binding-facilitated tissue uptake^40,41^.

Since the improved potency (on an ED_50_ basis) of the NMA SSO in *SMN2* splicing correction was consistent with a lower EC_50_, we hypothesized that the higher potency of the NMA SSO would be observed in cultured cells. Fibroblasts from an SMA patient were transfected by electroporation with increasing concentrations of the NMA-1 SSO or the MOE-1 SSO and *SMN2* splicing correction was analyzed after 24 h by real-time RT-PCR. The NMA-1 SSO was 1.6-fold more potent than the MOE-1 SSO (Extended Data Fig. 5, EC_50_ of NMA-1 = 0.18 µM vs MOE-1 = 0.28 µM). This suggests that the potency gain conferred by the NMA modification is the result of an intracellular interaction. However, this interaction is not an increased affinity for binding to that of the target site on the *SMN2* transcript since the Tm of the NMA-1 SSO is similar to the MOE-1 SSO (Supplementary information, Table S1).

### Optimizing the tolerability of the NMA-1 SSO

To evaluate the tolerability of the NMA-1 SSO we delivered a high dose to the CNS of wild-type mice by ICV bolus injection. Bolus injection of high dose SSOs into the mouse CSF can sometimes lead to various degrees of transient sedation, or death in the most severe case^42^. To evaluate the tolerability of the NMA-1 SSO we used a 7-point scale that we established previously where increasing scores reflect a more profound sedation 3 h after SSO administration^42^. ICV bolus injection of the NMA-1 SSO resulted in an acute sedation score of 5, manifested by the animal lying on its side without spontaneous movement after being transferred to a flat surface, while still responsive to a tail pinch. The NMA-1 SSO-mediated profound sedation was transient and the mice fully recovered after 24 h (Fig. 2A,B).

**Figure 2.**
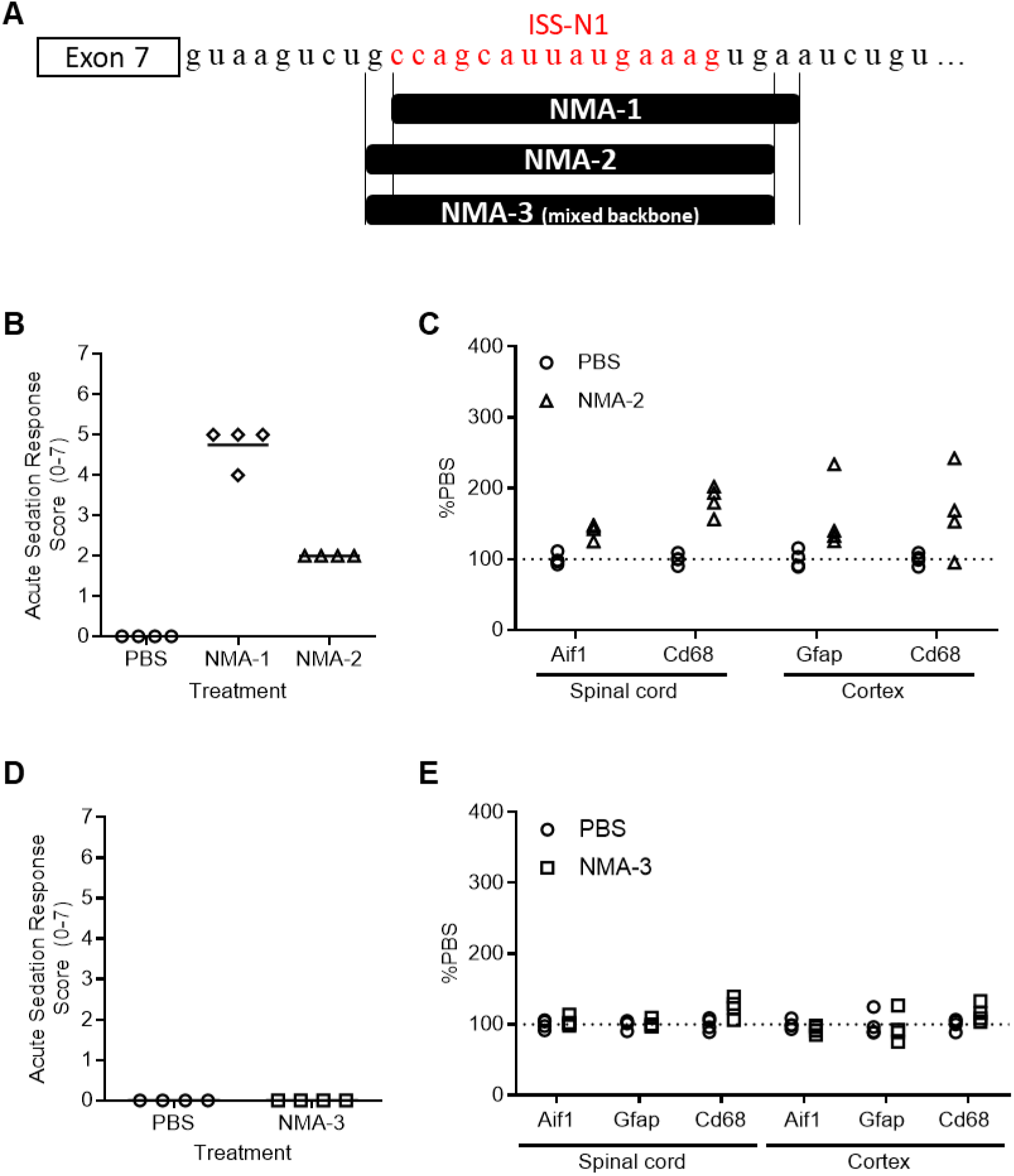
Sequence and chemistry optimization of NMA-1 SSO. (A) Schematic of positions of NMA SSOs on the *SMN2* pre-mRNA (partial sequence of intron 7 shown). (B) Acute sedation response scores in C57BL/6 mice after intracerebroventricular bolus injection of PBS, NMA-1 or NMA-2 SSOs. (C) Real-time RT-PCR of astrocyte marker gene *Gfap*, microglia marker gene *Aif1* and macrophage gene *Cd68* in spinal cord and cortex of C57BL/6 mice administered with PBS or NMA-2 SSO. (D) Acute sedation response scores in C57BL/6 mice after intracerebroventricular bolus injection of PBS or NMA-3 SSO. (E) Real-time RT-PCR of astrocyte marker gene *Gfap*, microglia marker gene *Aif1* and macrophage gene *Cd68* in spinal cord and cortex of C57BL/6 mice administered with PBS or NMA-3 SSO.

We have shown previously that the SSO sequence contributes significantly to the acute sedation^42^. To determine if the acute sedation caused by the NMA-1 SSO could be mitigated, we evaluated NMA-modified SSOs of different sequences that bound to sites within ISS-N1 at a different position than the NMA-1 SSO. An NMA-modified SSO (NMA-2) shifted by 1 bp (by loss of a 5’-T and gain of a 3’-C) relative to the NMA-1 SSO significantly improved the acute sedation (score of 2) after ICV bolus delivery to wild-type mice at a high dose (Fig. 2A,B).

We next evaluated the long-term tolerability of the NMA-2 SSO in the mouse CNS by measuring glial marker gene expression 8 weeks following ICV bolus injection. High dose ICV bolus injection of the NMA-2 SSO increased the expression of the glial markers (*Aif1*, *Gfap*, *Cd68*) which is indicative of neuroinflammation in the mouse CNS (Fig. 2C). In our experience, phosphorothioate (PS) linkages can influence long-term ASO tolerability^43–45^, we performed a systematic analysis of positional substitution of PS with phosphodiester (PO) linkages to assess their effects on tolerability. We found that strategic PS substitution with PO linkages at position 2 and 4 from the 5’ end of the SSO (as in SSO-22) significantly reduced neuroinflammation (Extended Data Fig. 6). We applied the same PO placement strategy to the shifted NMA-2 SSO and tested the SSO in wild-type mice. We confirmed that administration of this shifted NMA mixed backbone SSO (NMA-3) into the mouse CNS did not induce a neuroinflammatory response (Fig. 2E). Furthermore, it is known that reducing PS content in SSOs can mitigate acute tolerability^42^. This PO placement further improved acute tolerability relative to the full PS shifted NMA-2 SSO. (Fig. 2D). Altogether, the newly identified NMA-3 SSO has an excellent acute and delayed tolerability profile in mice.

### NMA-3 is more potent than nusinersen in human patient fibroblasts and in the CNS of human *SMN2* transgenic mice

Having identified a well-tolerated NMA SSO (NMA-3) we first assessed its activity in fibroblasts from a patient with SMA. Fibroblasts were transfected by electroporation with increasing concentrations of the NMA-3 SSO or nusinersen (MOE-1) and *SMN2* splicing correction was analyzed after 24 h by real-time RT-PCR. We observed a dose-dependent increase in *SMN2* splicing correction with the NMA-3 SSO, with an EC_50_ of 0.17 µM. The NMA-3 SSO was 1.6-fold more potent than the MOE-1 SSO that had an EC_50_ of 0.28 µM (Fig. 3A).

**Figure 3.**
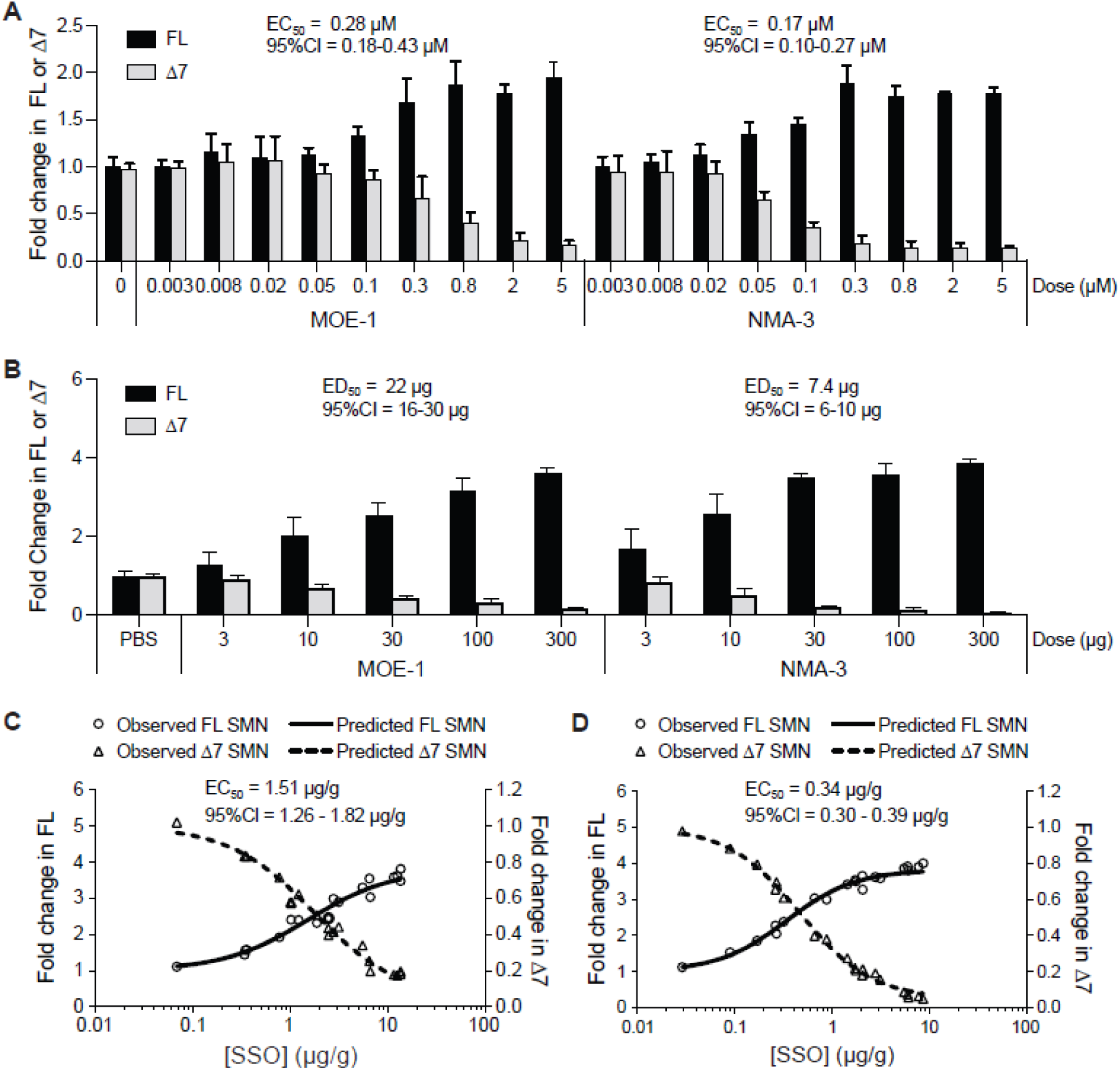
Potency for *SMN2* splicing correction of NMA-3 SSO compared with nusinersen (MOE-1). (A) Real-time RT-PCR analysis of *SMN2* transcripts including exon 7 (FL) or skipping exon 7 (Δ7) in SMA patient fibroblasts treated with MOE-1 or NMA-3 SSOs. For each dose level, n=3 duplicate culture wells. Error bars represent the S.D. The EC_50_ and confidence interval (CI) are shown. (B) Real-time RT-PCR analysis of FL or Δ7 *SMN2* transcripts in the spinal cord after intracerebroventricular (ICV) bolus injection of MOE-1 or NMA-3 SSOs in human *SMN2* transgenic mice. For each dose level, n=4. Error bars represent the S.D. The ED_50_ and CI are shown. (C) Concentration of MOE-1 SSO in the spinal cord plotted against the level of FL or Δ7 *SMN2* transcripts measured in the spinal cord of each mouse dosed with MOE-1 SSO in B. (D) Concentration of NMA-3 SSO in the spinal cord plotted against the level of FL or Δ7 *SMN2* transcripts measured in the spinal cord of each mouse dosed with NMA-3 SSO in B.

Next, we assessed the activity of the NMA-3 SSO in human *SMN2* transgenic mice. Increasing doses of the NMA-3 and MOE-1 SSOs were administered as a single ICV bolus injection and CNS tissues were analyzed for *SMN2* splicing 2 weeks after the injection by real-time RT-PCR. We observed a dose-dependent increase in *SMN2* splicing correction with the NMA-3 SSO with an ED_50_ in the spinal cord of 7.4 µg (Fig. 3B) and 17 µg in the brain (Extended Data Fig. 7A). The NMA-3 SSO was 3-fold more potent than the MOE-1 SSO. The MOE-1 SSO had an ED_50_ of 22 µg for exon 7 inclusion in the spinal cord (Fig. 3B) and 32 µg in the brain (Extended Data Fig. 7A). As expected, the NMA-3 SSO was also more potent than the MOE-1 SSO when the SSO levels in the CNS tissue were compared to the extent of *SMN2* splicing correction in each animal. The EC_50_ in the spinal cord was 0.34 µg/g for the NMA-3 SSO and was 1.51 µg/g for the MOE-1 SSO (Fig. 3C). The EC_50_ in the brain was 2.12 µg/g for the NMA-3 SSO and was 5.35 µg/g for the MOE-1 SSO (Extended Data Fig. 7B,C). Additionally, the improved potency of the NMA-3 SSO was attributable to the NMA modification and not its sequence or backbone composition (Extended Data Fig. 8).

### The NMA-3 SSO has a long duration of action in the CNS of human *SMN2* transgenic mice

Previously we demonstrated that in human *SMN2* transgenic mice nusinersen (MOE-1) has a long duration of action with sustained *SMN2* splicing correction for over 52 weeks after a single ICV bolus injection^34^. However, the NMA modification and PO modifications present in the NMA-3 SSO could have an impact on duration. Here we determined the duration of action for the NMA-3 SSO in human *SMN2* transgenic mice and compared it directly to the duration of action of the MOE-1 SSO. To account for the enhanced potency of the NMA-3 SSO we performed a single ICV bolus injection of 50 µg for the NMA-3 SSO and 100 µg for the MOE-1 SSO. The level of *SMN2* splicing correction was measured at various time points after the ICV bolus injection up to 52 weeks. The NMA-3 SSO displayed sustained *SMN2* splicing correction in the spinal cord and brain (Fig. 4, Extended Data Fig. 9), comparable to what was observed with the MOE-1 SSO. The long-lasting *SMN2* splicing correction observed with the NMA-3 SSO was attributable to its long half-life in the mouse CNS tissue (81 days in the spinal cord, and 196 days in the brain).

**Figure 4.**
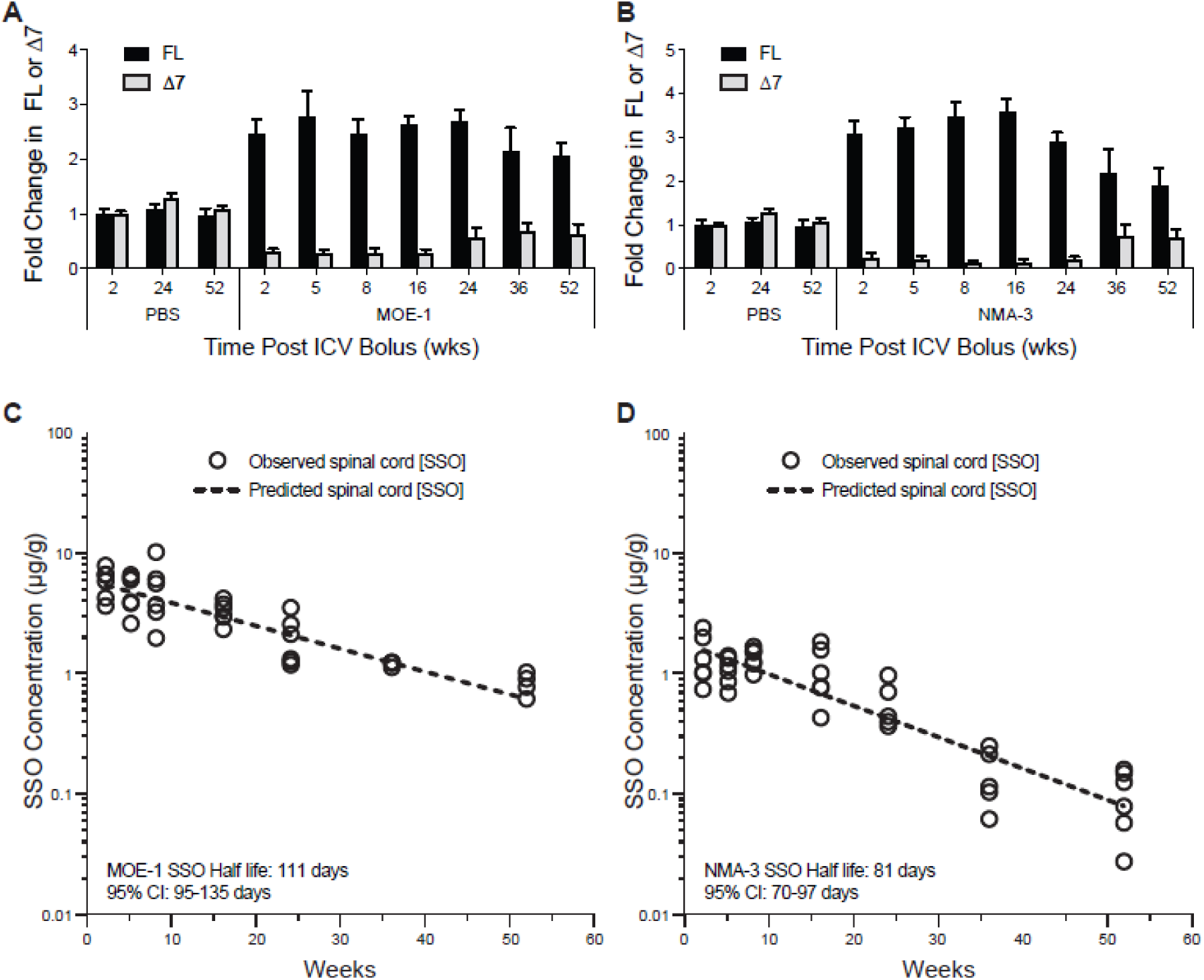
Duration of action and half-life of NMA-3 SSO compared with nusinersen (MOE-1) in the spinal cord of transgenic mice. (A) Real-time RT-PCR analysis of FL or Δ7 *SMN2* transcripts in the spinal cord at the indicated timepoints after intracerebroventricular (ICV) bolus injection of PBS or 100 µg of MOE-1 in the human *SMN2* transgenic mice. (B) Same as A except that 50 µg of NMA-3 SSOs were dosed instead of MOE-1. (C) Concentration of MOE-1 SSO in the spinal cord of each mouse dosed in A. Each open circle represents individual animal. (D) Concentration of NMA-3 SSO in the spinal cord of each mouse dosed in B.

### Enhanced potency of an NMA-modified *SCN1A* SSO

A therapeutic strategy that is being pursued for the treatment of Dravet syndrome is to use an SSO to promote skipping of exon 20N in transcripts produced from the *SCN1A* gene (Fig. 5A). This strategy has been shown to increase the levels of *SCN1A*, which is reduced in Dravet syndrome patients^26,27^. The SSO previously used to induce skipping of *SCN1A* exon 20N is an 18-mer uniformly modified with MOE referred to as STK-001^26,27^. To determine if the NMA modification could also be used to enhance splicing correction of *SCN1A*, we tested an NMA-modified SSO targeted to exon 20N with the same sequence as STK-001. We administered increasing doses of the NMA version of STK-001 (here called NMA-4) or STK-001 to wild-type mice via an ICV bolus injection. Brain tissue was analyzed for *SCN1A* transcripts with or without exon 20N by real-time RT-PCR 4 weeks post-injection. As expected, STK-001 showed a dose-dependent reduction in transcripts containing exon 20N, and an increase in transcripts that exclude exon 20N. The ED_50_ for splicing correction in the brain was 78 µg for STK-001 and the NMA-4 SSO was 3.5-fold more potent with an ED_50_ of 21 µg (Fig. 5B).

**Figure 5.**
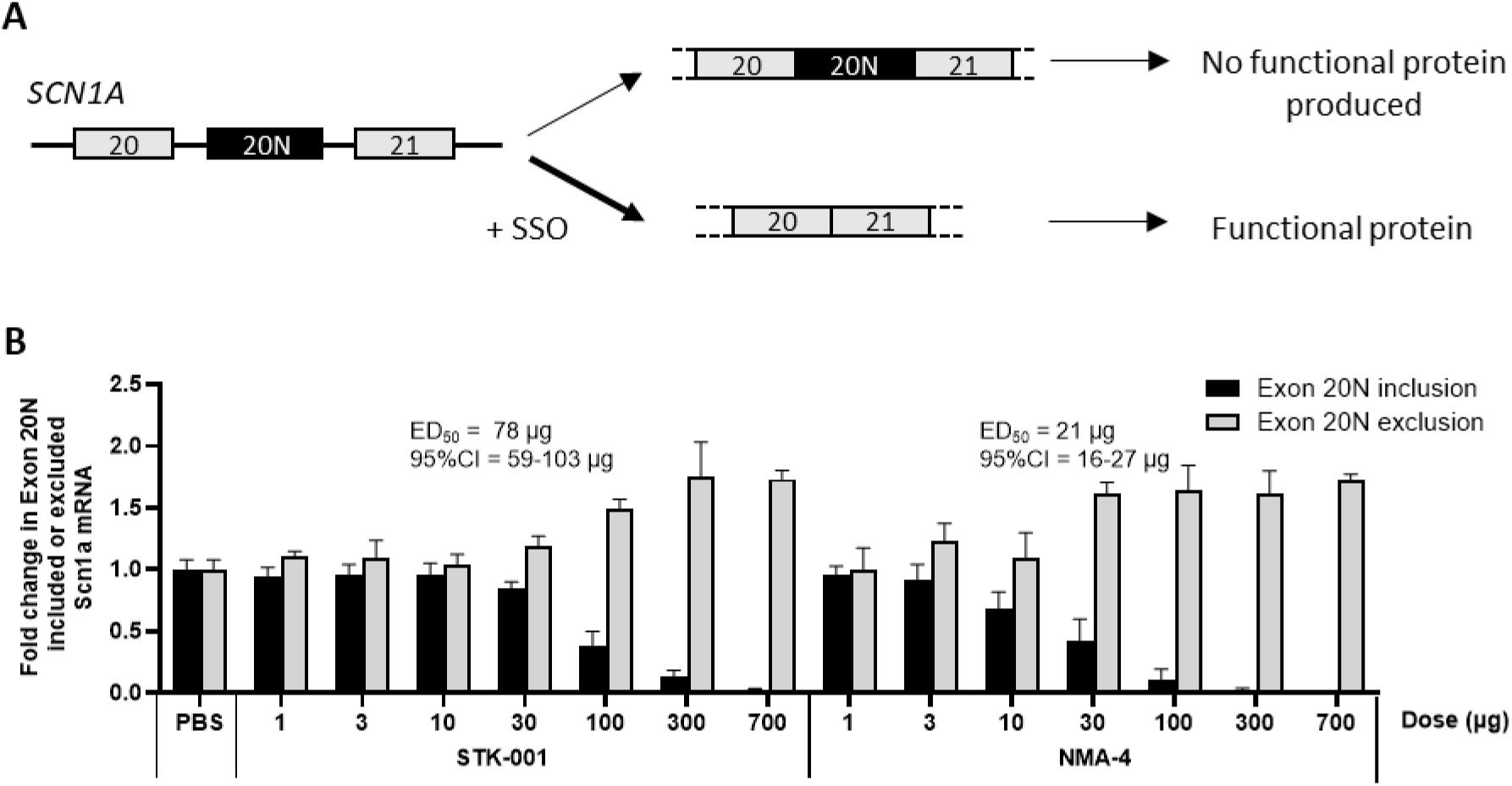
Potency comparison of MOE and NMA SSOs for splicing modulation of *SCN1A* mRNA in mice. (A) Schematic diagram of naturally occurring productive vs unproductive alternative splicing of *SCN1A* pre-mRNA. Exon 20N excluded transcript produces functional Nav1.1 protein whereas Exon 20N included transcript undergoes nonsense-mediated decay resulting in no functional protein production. Splice-modulating oligonucleotides (SSO) promotes exon 20N skipping to restore functional protein production. (B) Real-time RT-PCR analysis of mouse *Scn1a* transcripts including exon 20N (nonproductive transcript) or skipping exon 20N (productive transcript) in C57BL/6 mice after intracerebroventricular injection of different doses of STK-001 or NMA-4 SSOs. For each dose level, n=4 mice, except n=8 for PBS group and n=2 for STK-001 700 µg group. Error bars represent the S.D. The ED_50_ for exon 20N exclusion and confidence interval (CI) are shown.

## Discussion

Our work has identified a novel chemical modification, NMA, that significantly improves the activity of SSOs. NMA-modified SSOs are more potent than MOE-modified SSOs for correcting splicing of both *SMN2* exon 7 and *SCN1A* exon 20N. We believe that the NMA modification will also show higher potency for additional well established therapeutic targets where modulating splicing is required.

We demonstrated in fibroblasts from a patient with SMA and in human *SMN2* transgenic mice that an NMA modified SSO is significantly more potent for *SMN2* splicing correction than a MOE modified SSO. Additionally, we identified an NMA modified human candidate SSO (NMA-3) that is 3-4-fold more potent than nusinersen in our experimental models. Following extensive structure-activity relationship studies, we showed that NMA-3 is well tolerated in mice. Our newly identified NMA SSO also exhibits a long duration of action in human *SMN2* transgenic mice, which is similar to nusinersen. The current clinically approved dosing schedule for nusinersen is four loading doses in the first two months followed by dosing every four months. The higher potency of the NMA-3 SSO should allow for similar clinical efficacy to nusinersen but with less frequent dosing, thereby reducing the burden of repeated lumbar punctures for patients and caregivers. In fact, NMA-3 (salanersen) is currently being evaluated in a Phase 1 clinical trial (NCT05575011) in SMA patients with a once-yearly dosing regimen.

The exact mechanism by which the NMA modification enhances the potency of splicing modulation relative to the MOE modification is not known. What we currently know is that the NMA modification does not improve potency by providing a higher accumulation in CNS tissues since it accumulates to the same extent as the MOE-modified SSO (Fig. 3C,D). The fact that the NMA SSO has a lower EC_50_ for *SMN2* splicing correction in CNS tissues of human *SMN2* transgenic mice than the MOE SSO, and that the NMA SSO is more potent than the MOE SSO in cell culture when delivered by electroporation (Fig. 3A) suggests that the NMA modification is providing a benefit intracellularly. However, this is not due to an increased affinity for binding to its target site on the RNA since the Tm of NMA-modified SSOs is comparable to MOE-modified SSOs (Supplementary information, Table S1). Potential contributors to the improved activity conferred by NMA modification include improved release from endosomes^46,47^, better trafficking to the nucleus and/or the site of transcription on chromatin where splicing takes place^35,48^, or interactions with splicing factors or RNA binding proteins after the NMA SSO has hybridized to its target site as has previously been shown for 2’-F modified SSOs^35^. More work is necessary to decipher how the NMA chemistry improves the potency of SSOs.

## Methods

### Oligonucleotide synthesis

The synthesis and purification of the chemically modified splice-switching oligonucleotides (SSOs) were performed following the schemes described in the supplementary information. The oligonucleotides targeting human *SMN2* were 2’-*O*-MOE or 2’-*O*-NMA modified; SSOs were either 18 or 20 nucleotides in length. Parent SSO sequence (MOE-1) targeting the human *SMN2* gene was 5’-TCACTTTCATAATGCTGG-3’. SSO sequence targeting *SCN1A* gene was 5’-AGTTGGAGCAAGATTATC-3’. The lyophilized SSOs were dissolved and diluted to the desired concentration in sterile phosphate-buffered saline (PBS) without calcium or magnesium for experiments in mice. The SSO was quantified by UV spectrometry and sterilized by passage through a 0.2-µm filter before dosing.

### Intracerebroventricular administration of SSO in mice

Animal housing and all procedures met ethical standards for animal experimentation and were approved by the Institutional Animal Care and Use Committee at Ionis Pharmaceuticals. Adult male and female SMA Type III mice [FVB.Cg-*Smn1^tm1Hung^* Tg(SMN2)2Hung/J, stock number 005058] were obtained from The Jackson Laboratory (Bar Harbor, ME). Mice were anesthetized and placed on a stereotaxic frame fitted with a nose cone continuously supplied with 2% isoflurane. The scalp and anterior back were shaved and disinfected. A ∼1-cm incision was made on the scalp, and the subcutaneous tissues and periosteum were scraped from the skull with a sterile cotton-tipped applicator. A 10-μl Hamilton microsyringe with a 26 G Huber point removable needle was driven through the skull at 0.2 mm posterior and 1.0 mm lateral to bregma, and was lowered to a depth of 3 mm. Ten microliters of SSO solution or PBS were administered to the right cerebral ventricle at a rate of 1 µl per second. Three minutes after the injection, the needle was slowly withdrawn, and the incision was sutured. The mice were then allowed to recover from anesthesia in their home cage.

### RNA extraction and mRNA analysis

At necropsy, tissues collected were snap frozen at -70°C or below and later processed for RNA extraction and mRNA analysis by reverse-transcription real-time polymerase chain reaction (RT-qPCR). For mouse spinal cord, a 2-mm lumbar section was collected. For brain, a 1-mm coronal section, 2 mm posterior to the injection site was collected. RNA extraction and mRNA analyses were performed as previously described^34^ . Briefly, each piece of tissue was homogenized in a 2-ml tube containing Lysing Matrix D (MP Biomedicals, Santa Ana, CA), 500 µl of RLT buffer (Qiagen, Valencia, CA), and 1% (v/v) β-mercaptoethanol. Homogenization was performed for 20 seconds at 6000 rpm using a FastPrep Automated Homogenizer (MP Biomedicals). Ten microliters of lysate were used to isolate RNA with a RNeasy 96 Kit (Qiagen) that included in-column DNA digestion with 50U of DNase I (Invitrogen, Carlsbad, CA). Real-time RT-PCR was performed with the EXPRESS One-Step SuperScript qRT-PCR kit (Thermo Fisher Scientific) using gene-specific primers as described previously^34,35^. The FL or Δ7 *SMN2* expression level was normalized to that of *Gapdh* or total *SMN2* and further normalized to the level in vehicle (phosphate-buffered saline) treated animals or cells. For the analysis of glial fibrillary protein (*Gfap*), allograft inflammatory factor-1(*Aif1*), or *Cd68* expression, normalization was to the levels of *Gapdh* and further normalized to the level in vehicle (phosphate-buffered saline) treated animals. Primer sequences are listed in Supplementary information, Table S3.

### Quantification of SSO Tissue Concentration

A 1-mm mouse brain coronal section, 3 mm posterior to the injection site and the thoracic spinal cord, was collected for bioanalytical evaluation. Each piece of tissue was weighed, and the amount of SSO was then measured by various bioanalytical methods^49^ , including capillary gel electrophoresis coupled with UV detection, high-performance liquid chromatography (HPLC) coupled with UV detection (HPLC-UV), HPLC coupled with tandem mass spectrometry detection, or a hybridization-based enzyme-linked immunosorbent assay (HELISA). For HELISA, the probes have a sequence complementary to the SSO. The probe for MOE-1 contained biotin-tetraethylene glycol (TEG) at the 5’ end and digoxigenin at the 3’ end. The probe for NMA-1 contained digoxigenin at the 5’ end and biotin-TEG at the 3’ end.

### Data Analysis

Dose– and concentration–response relationships were analyzed using GraphPad Prism software version 6.0 or higher (GraphPad Software, San Diego, CA). Data were fit by nonlinear regression with a variable-slope (four-parameter) logistic model and normalized responses. For determination of half-maximal response values (ED₅₀ for dose–response and EC₅₀ for SSO concentration–response), minimal response values were constrained to 1 to improve curve fitting accuracy. ED_50_ and EC_50_ values were calculated from the fitted curves and are reported with 95% confidence intervals. The tissue half-life of SSOs associated with the apparent terminal elimination phase was calculated using a noncompartmental analysis (extravascular input model) applied to the mean concentration-time profile using Phoenix WinNonlin, Version 6.0 or higher.

---

**Extended Data Fig. 1.**
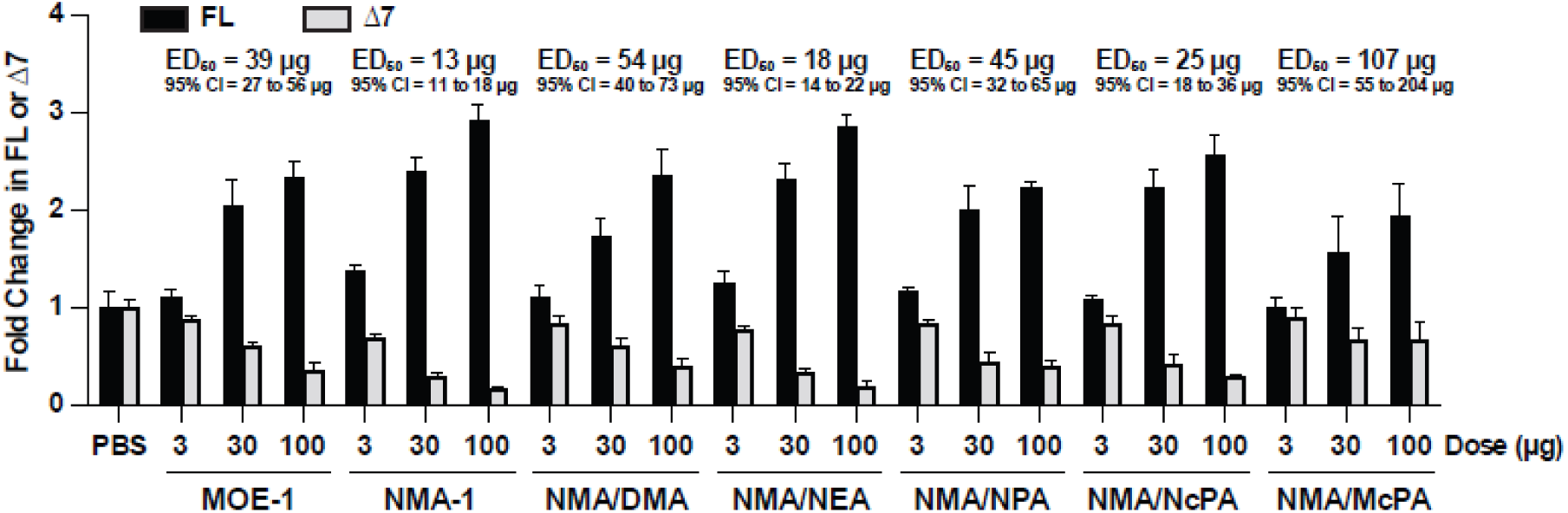
Potency for *SMN2* splicing correction of SSOs modified with MOE, NMA, or NMA analogs in the brain of the human *SMN2* transgenic mice. Real-time RT-PCR analysis of *SMN2* transcripts including exon 7 (FL) or skipping exon 7 (Δ7) in the brain 14 days after intracerebroventricular (ICV) bolus injection of MOE-1, NMA-1 or SSOs modified with analogs of NMA in *SMN2* transgenic mice. For each dose level, n=4. Error bars represent the S.D. The ED_50_ for exon 7 inclusion (FL) and confidence interval (CI) are shown.

**Extended Data Fig. 2.**
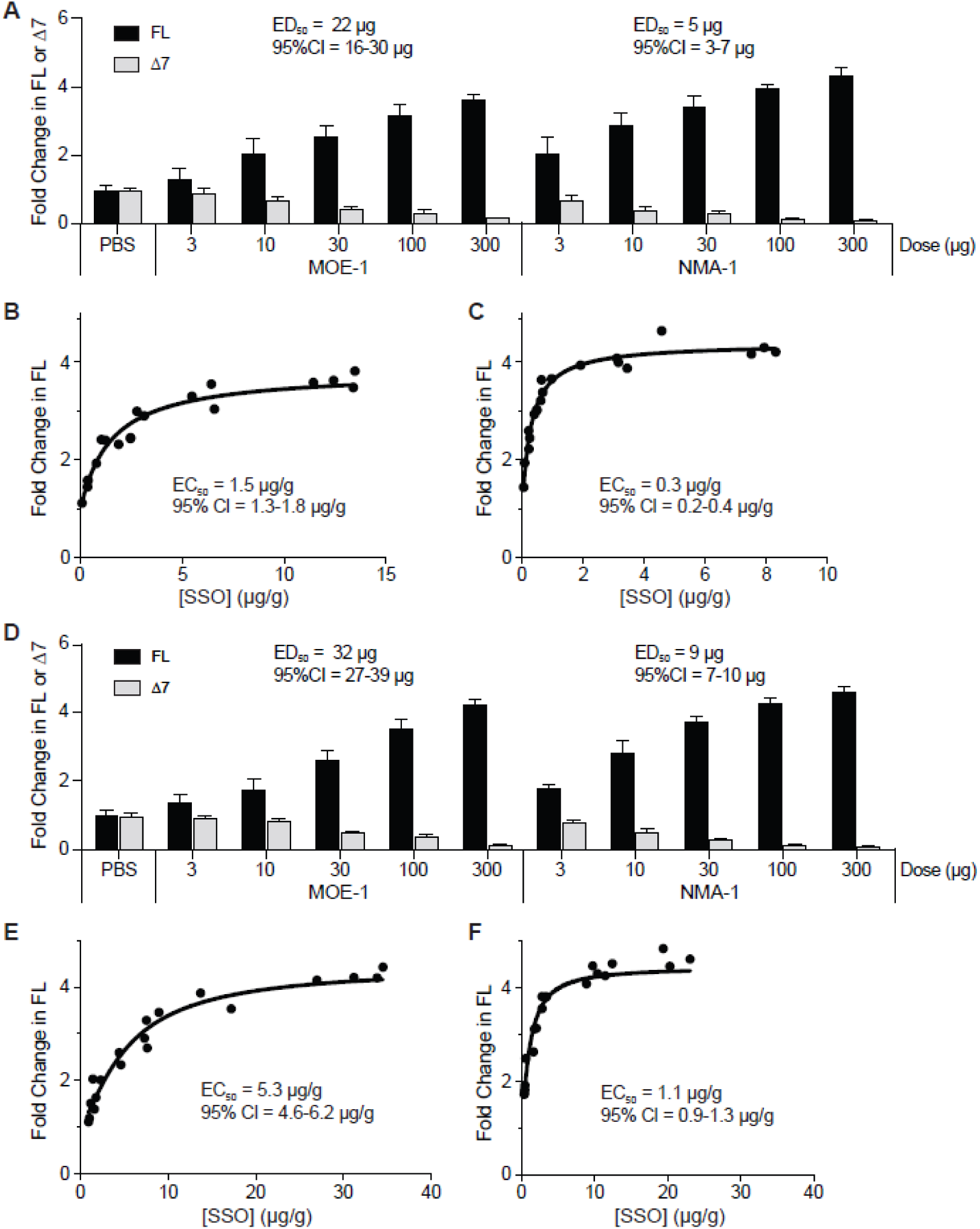
Potency for *SMN2* splicing correction of MOE-1 and NMA-1 SSOs in the spinal cord and brain of the human *SMN2* transgenic mice. (A). Real-time RT-PCR analysis of *SMN2* transcripts including exon 7 (FL) or skipping exon 7 (Δ7) in the spinal cord *SMN2* transgenic mice after ICV injection of MOE-1 or NMA-1 SSOs. For each dose level, n=4. The ED_50_ for exon 7 inclusion (FL)and confidence interval (CI) are shown. (B) Concentration of MOE-1 SSO plotted against the level of FL *SMN2* transcripts measured in the spinal cord of each mouse dosed with MOE-1 SSO. (C) Concentration of NMA-1 SSO plotted against the level of FL *SMN2* transcripts measured in the spinal cord of each mouse dosed with NMA-1 SSO in B. The EC_50_ for exon 7 inclusion (FL) and confidence interval (CI) are shown. (D) Same as A, except the real-time RT-PCR analysis was performed on brain tissues. (E) Concentration of MOE-1 SSO plotted against the level of FL *SMN2* transcripts measured in the brain of each mouse dosed with MOE-1 SSO. (F) Concentration of NMA-1 SSO plotted against the level of FL *SMN2* transcripts measured in the brain of each mouse dosed with NMA-1 SSO.

**Extended Data Fig. 3.**
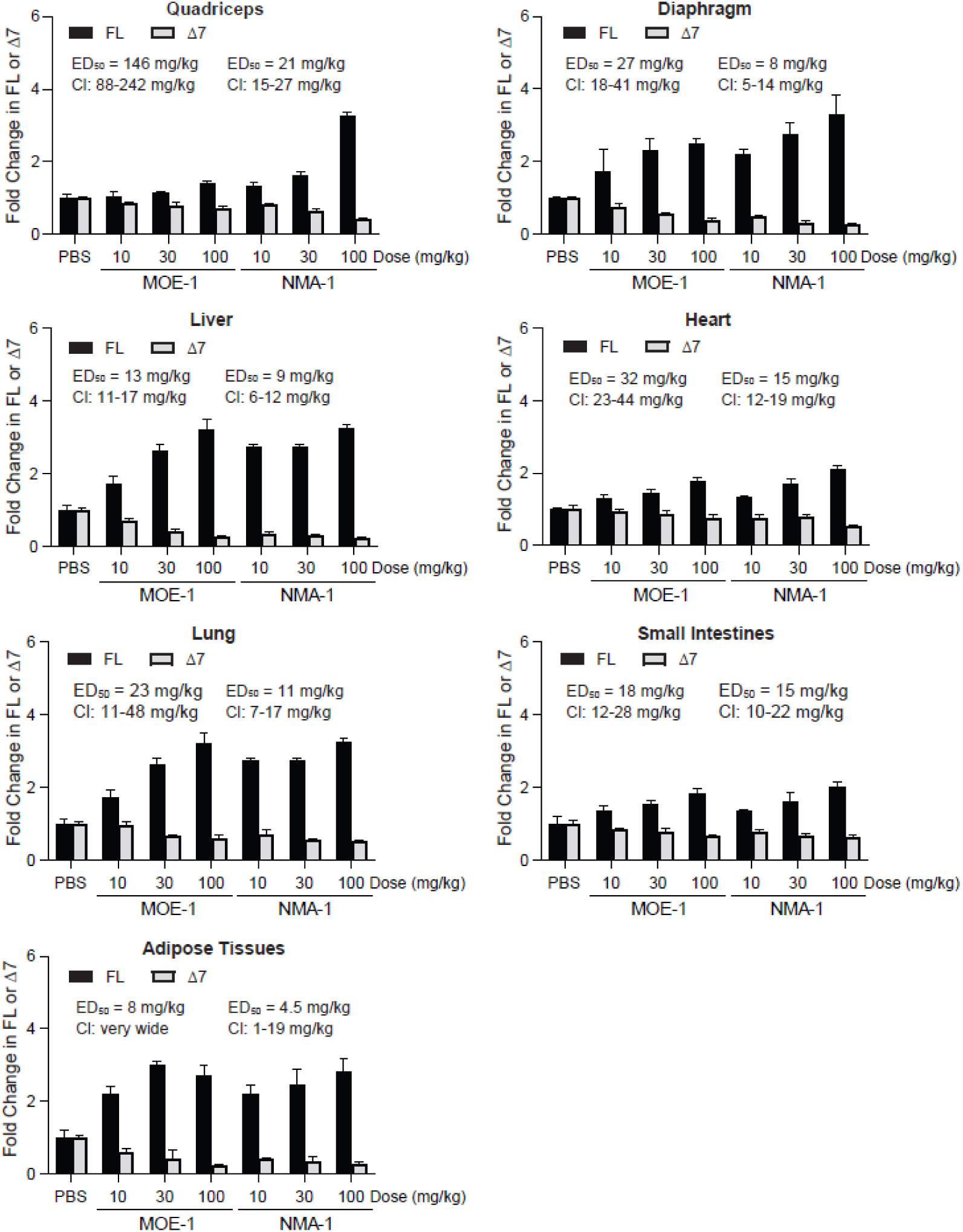
Potency for *SMN2* splicing correction of MOE-1 and NMA-1 in the peripheral tissues of the human *SMN2* transgenic mice. Real-time RT-PCR analysis of *SMN2* transcripts including exon 7 (FL) or skipping exon 7 (Δ7) in quadriceps, diaphragm, liver, heart, lung, small intestines and adipose tissues after subcutaneous injection with MOE-1 or NMA-1 in *SMN2* transgenic mice. For each dose group, n=4. Error bars represent the S.D. The ED_50_ for exon 7 inclusion (FL) and confidence interval (CI) are shown.

**Extended Data Fig. 4.**
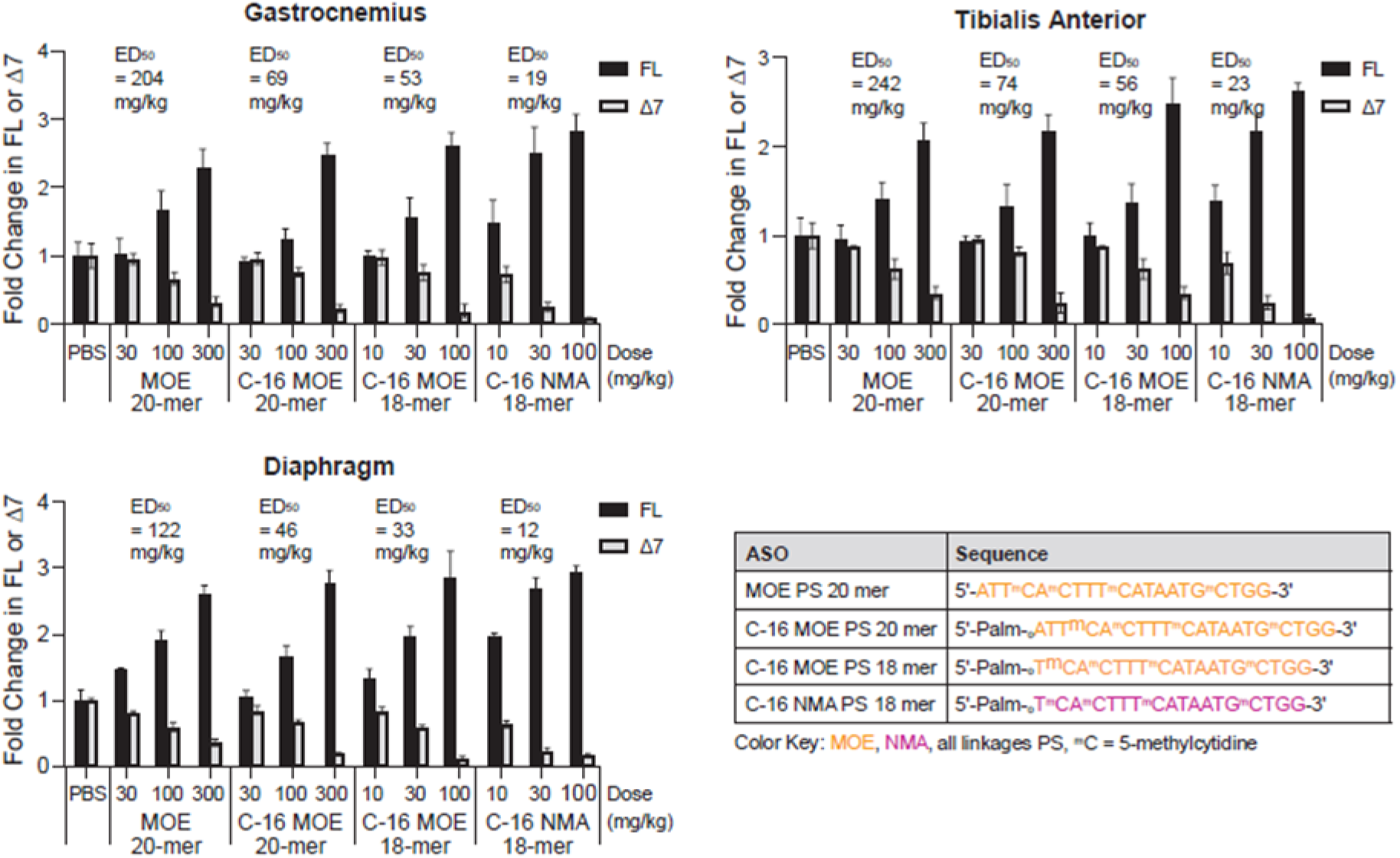
Potency comparison of C-16 conjugated MOE and NMA SSOs. (A) Real-time RT-PCR analysis of *SMN2* transcripts including exon 7 (FL) or skipping exon 7 (Δ7) in gastrocnemius, tibialis anterior and diaphragm after subcutaneous injection with unconjugated or C16-conjugated MOE or NMA SSOs in the human *SMN2* transgenic mice. For each dose group, n=4. Error bars represent the S.D. The ED_50_ for exon 7 inclusion (FL) is shown. Table shows the sequence and chemistry of each SSO.

**Extended Data Fig. 5.**
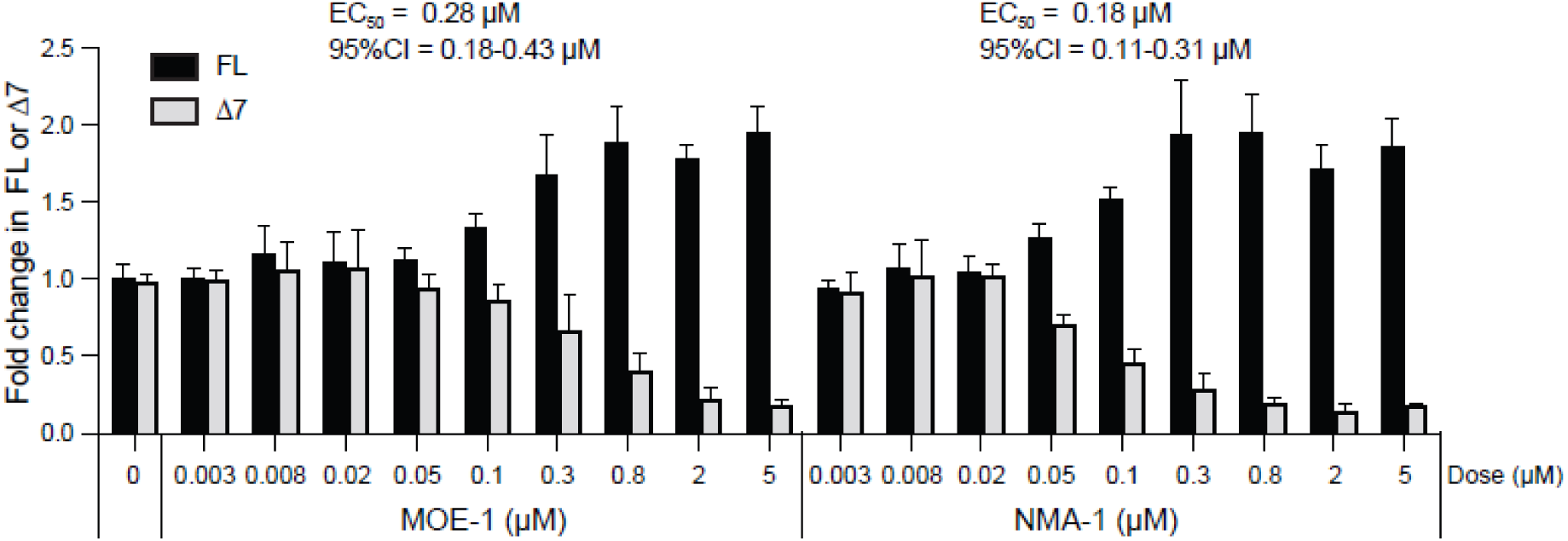
Potency for *SMN2* splicing correction of MOE-1 and NMA-1 in SMA patient fibroblasts. Real-time RT-PCR analysis of *SMN2* transcripts including exon 7 (FL) or skipping exon 7 (Δ7) in SMA patient fibroblasts treated with MOE-1 or NMA-1 SSOs. The ED_50_ for exon 7 inclusion (FL) and confidence interval (CI) are shown.

**Extended Data Fig. 6.**
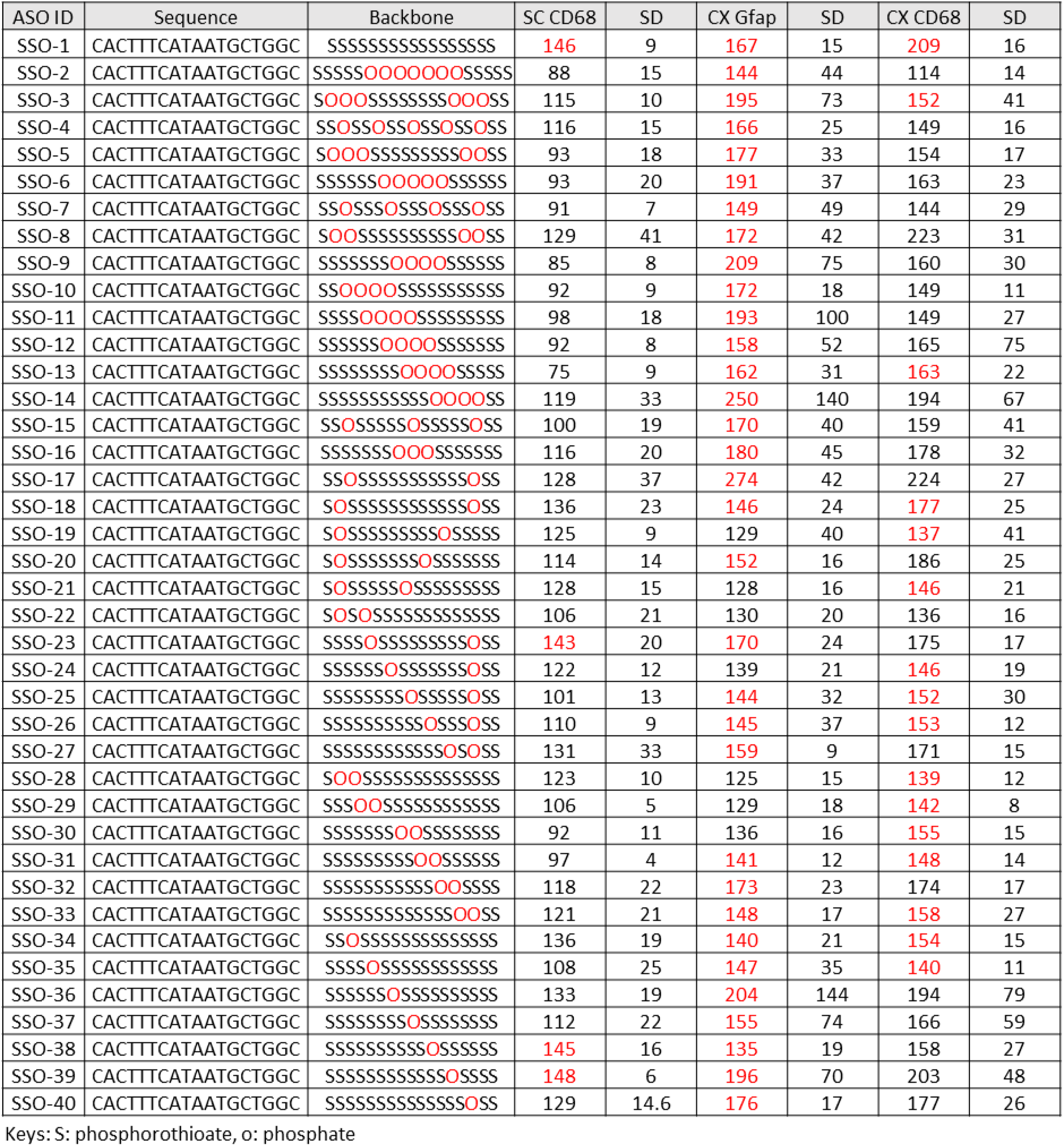
Effect of phosphate (PO) placement in backbone of the shifted MOE SSO on neuroinflammation. Real-time RT-PCR analysis of neuroinflammatory marker genes *Gfap* and *Cd68* in the cortex and the spinal cord of mice injected with the shifted MOE SSOs containing different designs of PO substitution in the backbone. SC: spinal cord. CX: cortex. Values represent mRNA levels relative to PBS (vehicle) control in %. SD is standard deviation.

**Extended Data Fig. 7.**
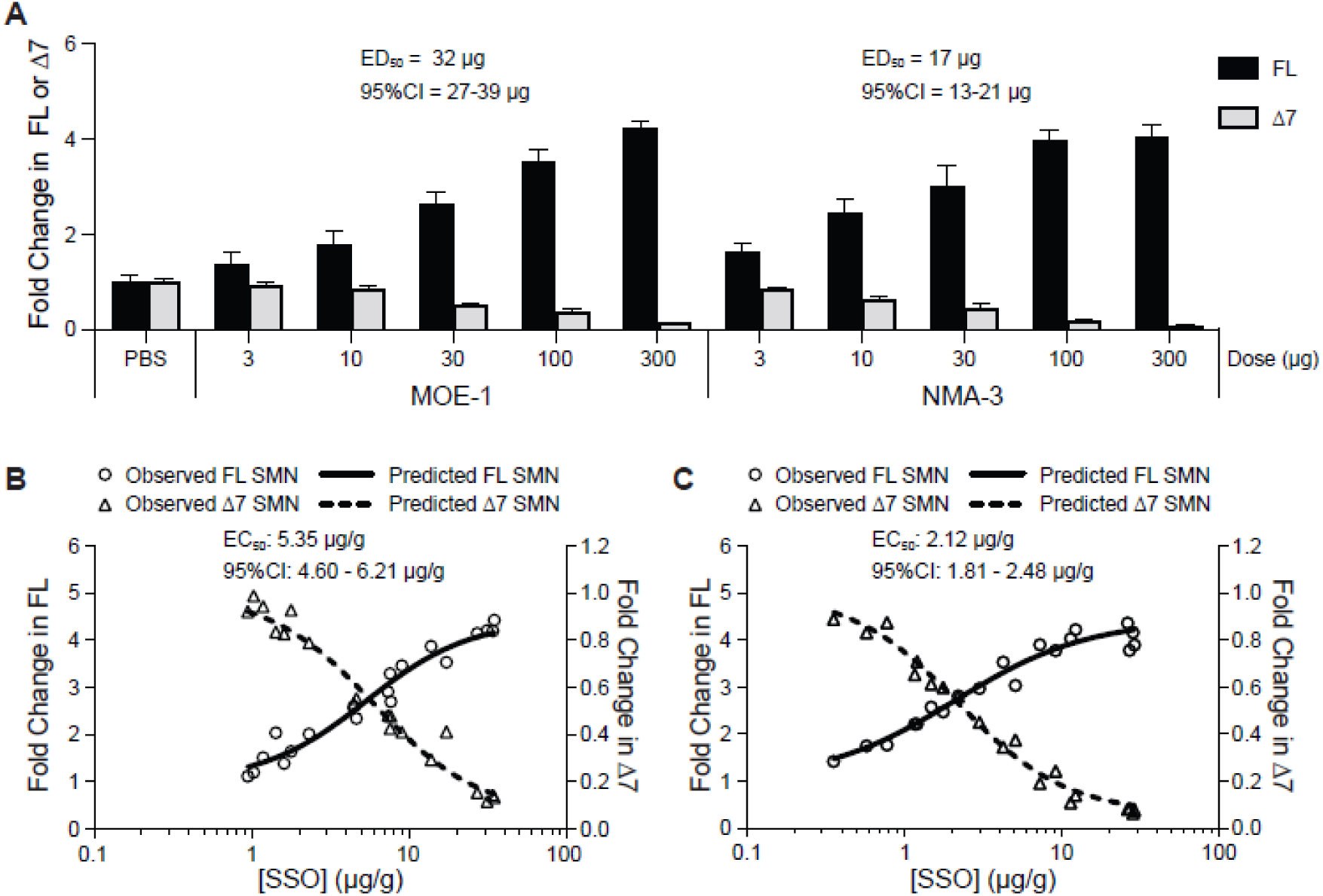
Potency for *SMN2* splicing correction of NMA-3 SSO compared with nusinersen (MOE-1) in the brain of human *SMN2* transgenic mice. (A) Real-time RT-PCR analysis of FL or Δ7 *SMN2* transcripts in the brain 14 days after intracerebroventricular (ICV) bolus injection of MOE-1 or NMA-3 SSOs in the *SMN2* mice. For each dose level, n=4. Error bars represent the S.D. The ED_50_ for exon 7 inclusion (FL) and confidence interval (CI) are shown. (B) Concentration of MOE-1 SSO plotted against the level of FL or Δ7 *SMN2* transcripts measured in the brain of each mouse dosed with MOE-1 SSO in A. (C) Concentration of NMA-3 SSO plotted against the level of FL or Δ7 *SMN2* transcripts measured in the brain of each mouse dosed with NMA-3 SSO in A. Each data point represents an individual animal. The EC_50_ for exon 7 inclusion (FL) and confidence interval (CI) are shown.

**Extended Data Fig. 8.**
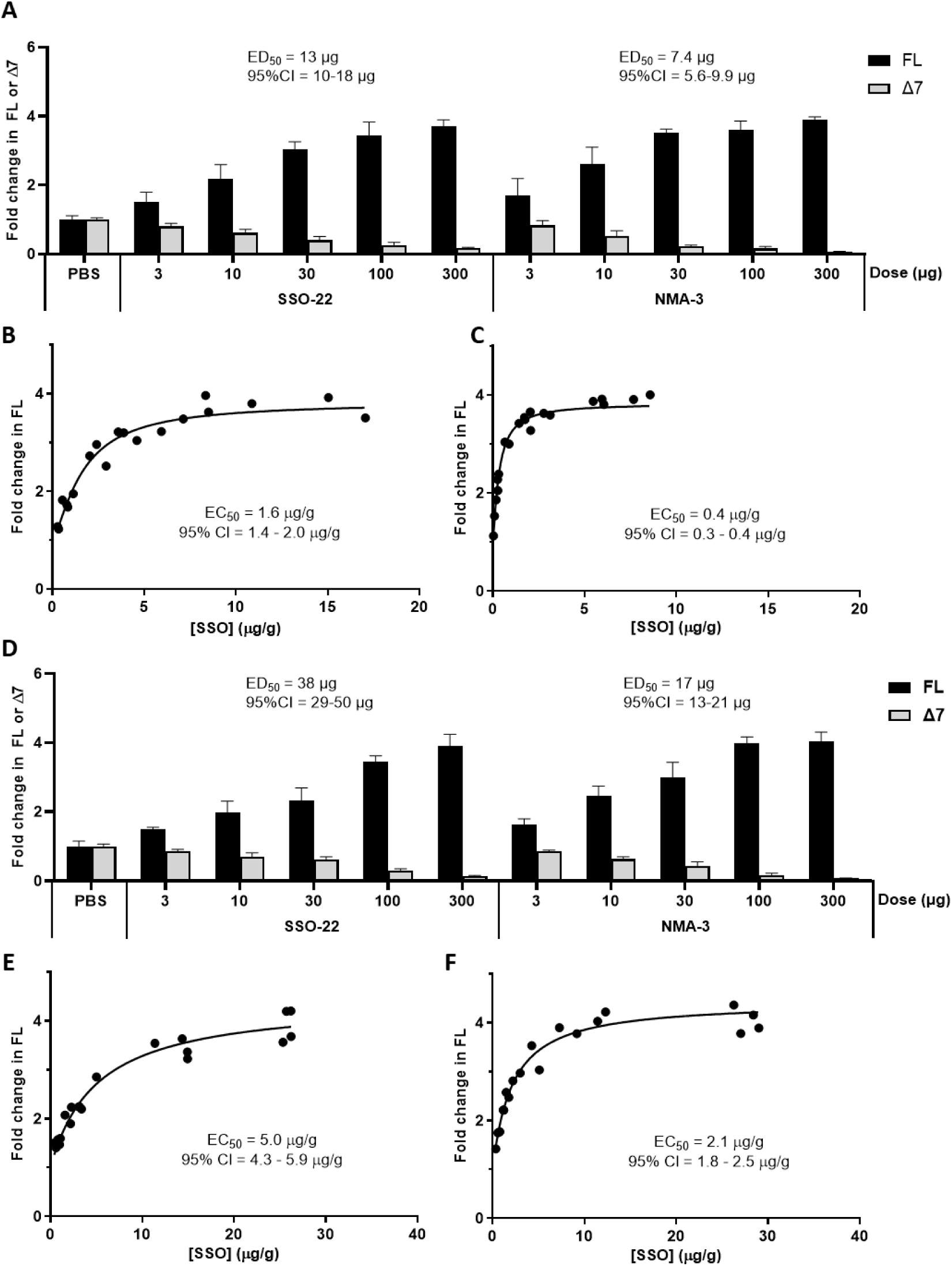
Potency for *SMN2* splicing correction of NMA-3 SSO compared with MOE SSO with the same sequence and backbone composition (SSO-22). (A) Real-time RT-PCR analysis of *SMN2* transcripts including exon 7 (FL) or skipping exon 7 (Δ7) in the spinal cord *SMN2* transgenic mice at 14 days after ICV injection with SSO-22 or NMA-3 SSOs. For each dose level, n=4. The ED_50_ for exon 7 inclusion (FL) and confidence interval (CI) are shown. (B) Concentration of SSO-22 SSO plotted against the level of FL *SMN2* transcripts measured in the spinal cord of each mouse dosed with SSO-22 SSO. (C) Concentration of NMA-3 SSO plotted against the level of FL *SMN2* transcripts measured in the spinal cord of each mouse dosed with NMA-3 SSO. (D) Same as A, except the real-time RT-PCR analysis was performed on brain tissues. (E) Concentration of SSO-22 SSO plotted against the level of FL *SMN2* transcripts measured in the brain of each mouse dosed with SSO-22 SSO. (F) Concentration of NMA-3 SSO plotted against the level of FL *SMN2* transcripts measured in the brain of each mouse dosed with NMA-3 SSO.

**Extended Data Fig. 9.**
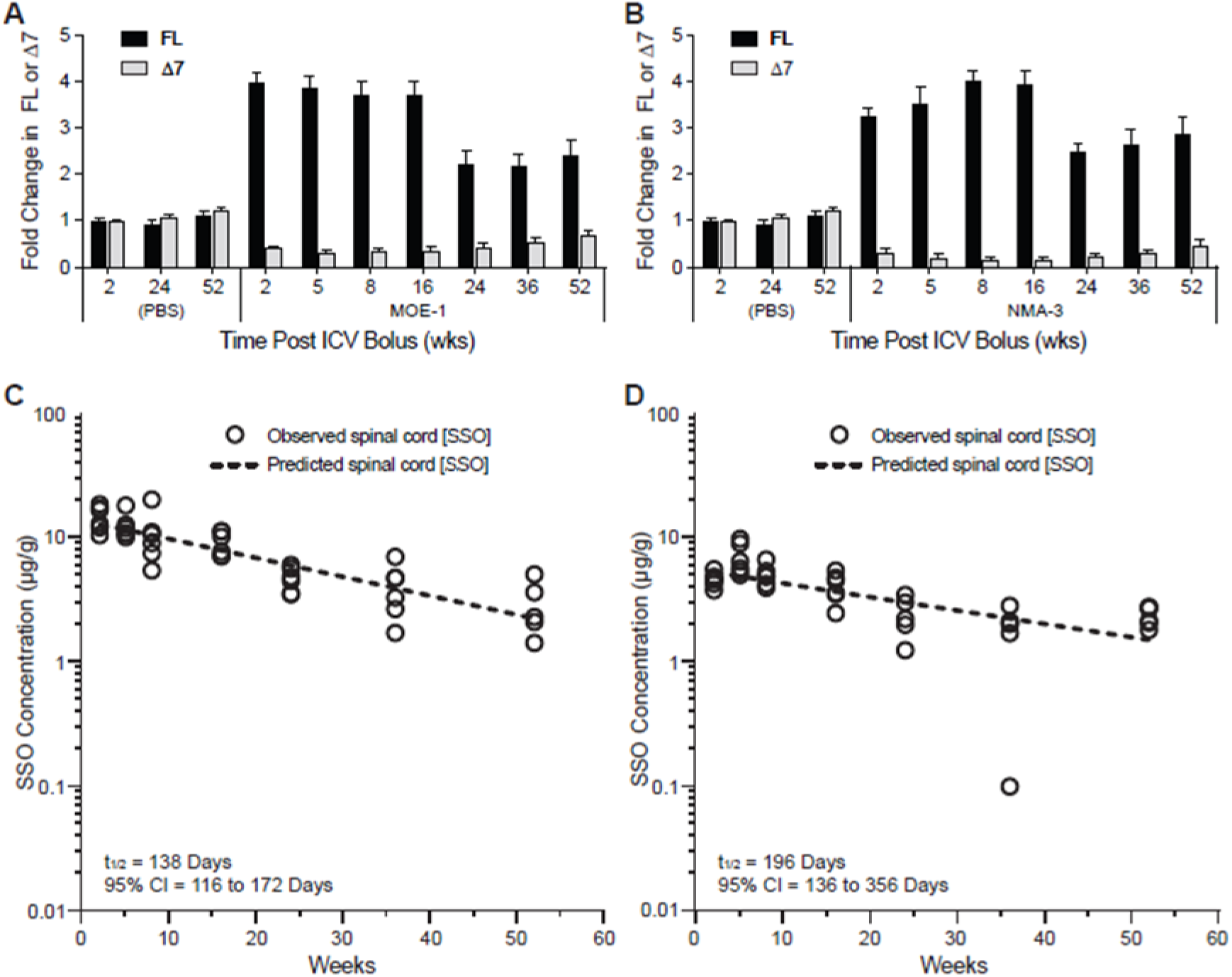
Duration of action and half-life of NMA-3 compared with nusinersen (MOE-1) in the brain of transgenic mice. (A) Real-time RT-PCR analysis of *SMN2* transcripts including exon 7 (FL) or skipping exon 7 (Δ7) in the brain of human *SMN2* transgenic mice injected with MOE-1 SSO. (B) Same as A except that the NMA-3 SSOs were dosed instead of MOE-1. (C) The amount of MOE-1 SSO in the brain of each mouse was measured at 2 to 16-week timepoint by HPLC-UV and 24-54-week timepoint by HELISA. (D) The amount of NMA-3 measured in the brain of each mouse. Each open circle represents individual animal.

