## Supplementary information for "Enhanced splicing modulation by NMA-modified antisense oligonucleotides"

*Ionis Pharmaceuticals, 2855 Gazelle Ct., Carlsbad, CA 92010, USA*

**Table S1.** Sequence, chemistry, and Tm of ASOs containing NMA and its analogs

| ASO ID | Sequence and Chemistry | $T_m$ °C | Δ $T_m$ °C |
|---|---|---|---|
| MOE-1 | 5'-T$^m$CA$^m$CTTT$^m$CATAATG$^m$CTGG-3' | 68.1 | - |
| NMA-1 | 5'-T$^m$CA$^m$CTTT$^m$CATAATG$^m$CTGG-3' | 70.7 | 2.6 |
| NMA/DMA | 5'-T$^m$CA$^m$CTTT$^m$CATAATG$^m$CTGG-3' | 60.2 | -7.9 |
| NMA/NEA | 5'-T$^m$CA$^m$CTTT$^m$CATAATG$^m$CTGG-3' | 71.1 | 3 |
| NMA/NPA | 5'-T$^m$CA$^m$CTTT$^m$CATAATG$^m$CTGG-3' | 70.4 | 2.3 |
| NMA/NcPA | 5'-T$^m$CA$^m$CTTT$^m$CATAATG$^m$CTGG-3' | 70.9 | 2.8 |
| NMA/McPA | 5'-T$^m$CA$^m$CTTT$^m$CATAATG$^m$CTGG-3' | 71.6 | 3.5 |

Color Key: MOE, NMA, NMA analogs, all linkages PS, $^m$C = 5-methylcytidine

**Table S2.** Analytical data for ASOs containing NMA and its analogs

| ASO ID | Calcd Mass | Found Mass | %UV Purity |
|---|---|---|---|
| MOE-1 | 7,127.32 | 7126.12 | 95.01 |
| NMA-1 | 7,361.2 | 7360.04 | 95.95 |
| NMA/DM | 7515.45 | 7516.16 | 87.95 |
| NMA/NEA | 7515.45 | 7514.68 | 87.70 |
| NMA/NPA | 7669.74 | 7669.16 | 89.19 |
| NMA/NcP | 7647.26 | 7646.96 | 88.69 |
| NMA/McP | 7801.68 | 7801.08 | 89.25 |

**Table S3.** Primer sequences used in this study

| Transcript | Primer/probe | Sequence |
|---|---|---|
| Total *SMN2* | Forward | CAGGAGGATTCCGTGCTGTT |
| | Reverse | CAGTGCTGTATCATCCCAAATGTC |
| | Probe | ACAGGCCAGAGCGAT |
| *SMN2* FL | Forward | GCTGATGCTTTGGGAAGTATGTTA |
| | Reverse | CACCTTCCTTCTTTTTGATTTTGTC |
| | Probe | TACATGAGTGGCTATCATACT |
| *SMN2* Δ7 | Forward | CATGGTACATGAGTGGCTATCATACTG |
| | Reverse | TGGTGTCATTTAGTGCTGCTCTATG |
| | Probe | CCAGCATTTCCATATAATAGC |
| *Aif1* | Forward | TGGTCCCCCAGCCAAGA |
| | Reverse | CCCACCGTGTGACATCCA |
| | Probe | AGCTATCTCCGAGCTGCCCTGATTGG |
| *Cd68* | Forward | TGGCGGTGGAATACAATGTG |
| | Reverse | GATGAATTCTGCGCCATGAA |
| | Probe | CCTTCCCACAGGCAGCACAGTGG |
| *Gfap* | Forward | GAGAGAGATTCGCACTCAATACGA |
| | Reverse | GTCTGCAAACTTAGACCGATACCA |
| | Probe | CAGTGGCCACCAGTAACATGCAAGAGAC |
| *Gapdh* | Forward | GGCAAATTCAACGGCACAGT |
| | Reverse | GGGTCTCGCTCCTGGAAGAT |
| | Probe | AAGGCCGAGAATGGGAAGCTTGTCATC |
| *SCN1A* Exon 20X included | Forward | AGCCCTTTATTATGGGTGGTT |
| | Reverse | CCAGAATATAAGGCAAACCAGAAG |
| | Probe | TGGATGGAATTGCTCCTAACAGGGC |
| *SCN1A* Exon 20X excluded | Forward | CCCTAAGAGCCTTATCACGATTT |
| | Reverse | GGCAAACCAGAAGCACATTC |
| | Probe | AGGGTGGTTGTGAATGCCCTGTTA |

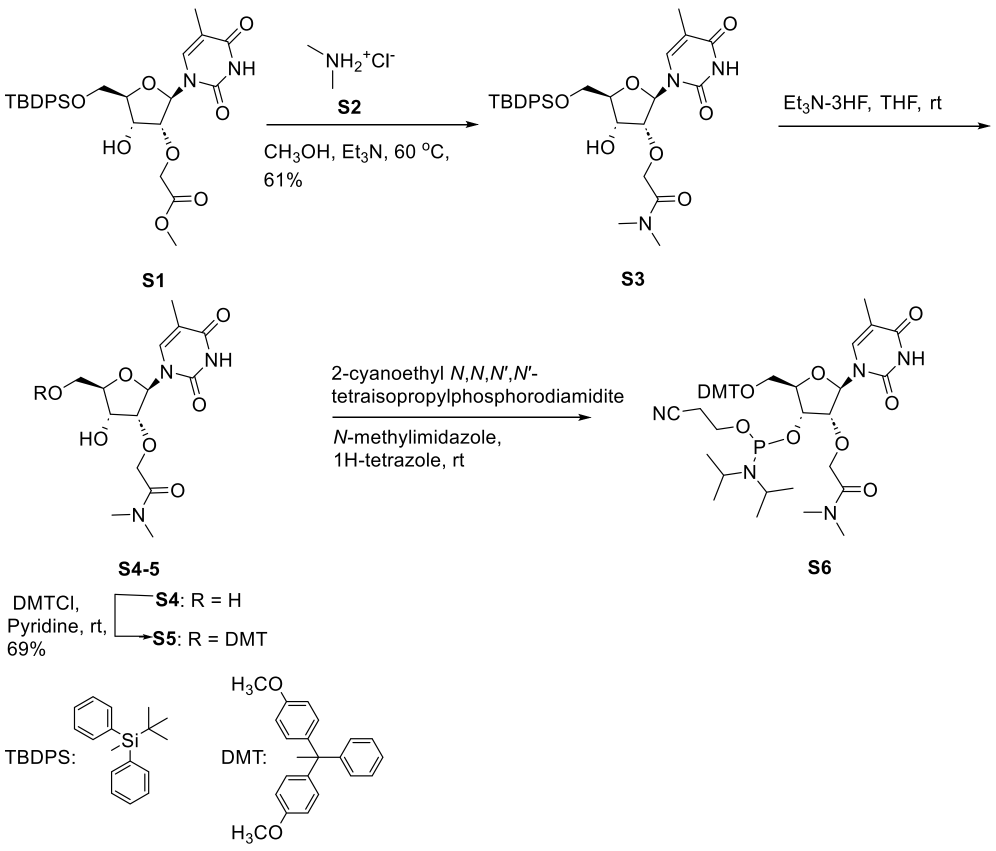

**Scheme S1**. Synthesis of 5'-*O*-DMT-2'-*O*-(DMA)-thymidine-3'-phosphoramidite **S6**

**Compound S3.** To a solution of compound **S1** (22.0 g, 37.8 mmol) in methanol (100 mL), dimethylamine hydrochloride **S2** (46.6 g, 566.0 mmol) and TEA (100.0 mL, 171.0 mmol) were added. After stirring the reaction mixture at 80 °C for 12 h in a pressure flask, it was concentrated to dryness. The residue was redissolved in dichloromethane (100 mL), washed with water (100 mL), and concentrated under reduced pressure. The residue was purified by a silica gel column chromatography and eluted first with 20% ethyl acetate in dichloromethane and then with 5% MeOH in dichloromethane to give **S3** (13.3 g, 60.7%). $^1$H NMR (300 MHz, $CDCl_3$): δ 9.38 (s, 1H), 7.62-7.75 (m, 4H), 7.58 (s, 1H), 7.31-7.48 (m, 6H), 5.95 (d, *J* = 2.6 Hz, 1H), 5.82 (d, *J* = 4.0 Hz, 1H), 4.38-4.66 (m, 2H), 4.26-4.36 (m, 1H), 4.10-4.22 (m, 2H), 3.89-4.01 (m, 2H), 2.98 (s,

3H), 2.92 (s, 3H), 1.47 (s, 3H), 1.10 (s, 9H); LRMS (ESI TOF): *m/z* calcd for $C_{30}H_{39}N_3NaO_7Si$: 604.3 , found 604.3 $[M + Na]^+$.

**Compound S4**. To a solution of compound **S3** (13.1 g, 22.5 mmol) in THF (150 mL) triethylamine (9.83 mL, 70.5 mmol)) and triethylamine trihydrofluoride (11.0 mL, 67.6 mmol) were added. The reaction mixture was concentrated after stirring for 2 h at room temperature. The residue obtained was purified by silica gel column chromatography and eluted with 0-30% MeOH in THF to yield **S4 (**8.3 g, quantitative). $^1$H NMR (300 MHz, DMSO-$d_6$): δ 11.32 (br s, 1H), 7.73 (s, 1H), 5.90 (d, *J* = 6.0 Hz, 1H), 5.70 (d, *J* = 3.1 Hz, 1H), 5.12 (t, *J* = 4.9 Hz, 1H), 4.20-4.51 (m, 2H), 3.98-4.10 (m, 2H), 3.81-3.93 (m, 1H), 3.48-3.70 (m, 2H), 2.84-2.91 (m, 3H), 2.82 (s, 3H), 1.77 (m, 3H) ; $^{13}$C NMR (75 MHz, DMSO-$d_6$): δ 169.7, 163.7, 150.7, 136.2, 109.4, 85.8, 85.1, 82.9, 68.7, 68.5, 60.9, 35.3, 35.0, 12.2; LRMS (ESI TOF): *m/z* calcd for $C_{14}H_{21}N_3NaO_7$: 366.1, found 366.1 $[M + Na]^+$.

**Compound S5**. Compound **S4** (8.34 g, 24.3 mmol), was dissolved in anhydrous pyridine (250 mL) and DMTCl (12.4 g, 36.5 mmol) was added at 0 $^o$C. The reaction was stirred at room temperature for 12 h and quenched by adding methanol (5 mL). The mixture was stirred at room temperature for 30 minutes and extracted with ethyl acetate (500 mL). The ethyl acetate solution was washed with water (500 mL) and concentrated to dryness under reduced pressure. The residue was purified by silica gel column chromatography and eluted first with 0-50% ethyl acetate in the hexanes (3-4 column volume) followed with 0-5% MeOH in dichloromethane to give **S5** (13.3 g, 85%).$^1$H NMR (300 MHz, $CDCl_3$): δ 9.30 (s, 1H), 7.72 (s, 1H), 7.39-7.46 (m, 2H), 7.17-7.36 (m, 7H), 6.71-6.95 (m, 4H), 5.86 (d, *J* = 1.7 Hz, 2H), 4.56 (q, *J* = 16.1 Hz, 2H), 4.38-4.47 (m, 1H), 4.16-4.24 (m, 1H), 3.98 (dd, *J* = 1.5, 5.2 Hz, 1H), 3.78 (s, 6H), 3.41-3.61 (m, 2H), 2.98 (s, 3H), 2.93 (s, 3H), 1.35 (s, 3H); $^{13}$C NMR (75 MHz, $CDCl_3$): δ 170.2, 163.9, 158.6, 158.6, 150.5, 144.5, 135.6, 135.5, 135.2, 130.1, 128.2, 127.9, 127.0, 113.3, 110.6, 89.3, 86.6, 86.0, 83.7, 68.8, 68.0,

61.7, 55.2, 35.7, 35.5, 11.8; LRMS (ESI TOF): *m/z* calcd for $C_{35}H_{39}N_3NaO_9$: 668.3, found 668.3 [M + Na]$^+$.

**5’-*O*-DMT-2’-*O*-(DMA)-thymidine-3’-phosphoramidite S6**. To a solution of compound **S5** (9.1 g, 14.0 mmol) DMF (70 mL) tetrazole (0.8 g, 11.2 mmol), *N*-methylimidazole (0.3 mL, 3.52 mmol) and 2-cyanoethyl *N*,*N*,*N′*,*N′*-tetraisopropylphosphorodiamidite (6.7 mL, 21.0 mmol) were added at 0 $^{o}$C (ice bath). The reaction mixture was removed from the ice bath and stirred for 2 h at room temperature. Diluted the reaction mixture with ethyl acetate (150 mL) and washed with saturated aqueous sodium bicarbonate solution (250 mL), brine (250 mL), dried ($Na_2$SO4), filtered, and concentrated under reduced pressure. The residue product was purified by silica gel column chromatography and eluted with 20-50% acetone in dichloromethane to yield **S6** (12.6 g, <95%). $^{31}$P NMR (121MHz, $CDCl_3$): δ 150.5; HRMS (ESI TOF): *m/z* calcd for $C_{44}H_{55}N_5O_{10}P$: 845.3765, found 844.3679 [M - H]$^-$.

**Compound S8**. Compound **S1** (20.3 g, 34.9 mmol) was dissolved in methanol (100 mL), and ethylamine solution **S7** (100.0 mL, 2M) and TEA (19.4 mL, 139.0 mmol) were added. The reaction mixture was stirred 50 $^{o}$C for 12 h and concentrated under reduced pressure. The residue was purified by silica gel column chromatography and eluted first with 30-40% ethyl acetate in a dichloromethane and then with 5-8% MeOH in dichloromethane to give **S8** (14.9 g, 74%).$^{1}$H NMR (300 MHz, $CDCl_3$): δ 10.27 (br s, 1H), 7.67 (t, *J* = 6.0 Hz, 4H), 7.65 (s, 1H), 7.33-7.51 (m, 6H), 7.12 (t, *J* = 5.4 Hz, 1H), 6.67 (br s, 1H), 5.96 (d, *J* = 3.1 Hz, 1H), 4.65 (d, *J* = 3.6 Hz, 1H), 4.21-4.43 (m, 3H), 4.11-4.20 (m, 1H), 3.86-4.00 (m, 2H), 3.19-3.43 (m, 2H), 1.49 (d, *J* = 0.9 Hz, 3H), 1.16 (t, *J* = 6.0 Hz, 3H), 0.99-1.11 (m, 9H); $^{13}$C NMR (75 MHz, $CDCl_3$): δ 171.7, 169.6, 163.9, 151.3, 135.5, 135.2, 134.7, 133.1, 132.3, 130.2, 128.0, 111.4, 88.2, 84.7, 84.3, 69.9, 68.0,

62.6, 62.1, 34.2, 33.9, 27.0, 19.4, 14.7, 14.4, 11.9; LRMS (ESI TOF): *m/z* calcd for $C_{30}H_{39}N_3NaO_7Si$: 604.3 , found 604.3 $[M + Na]^+$.

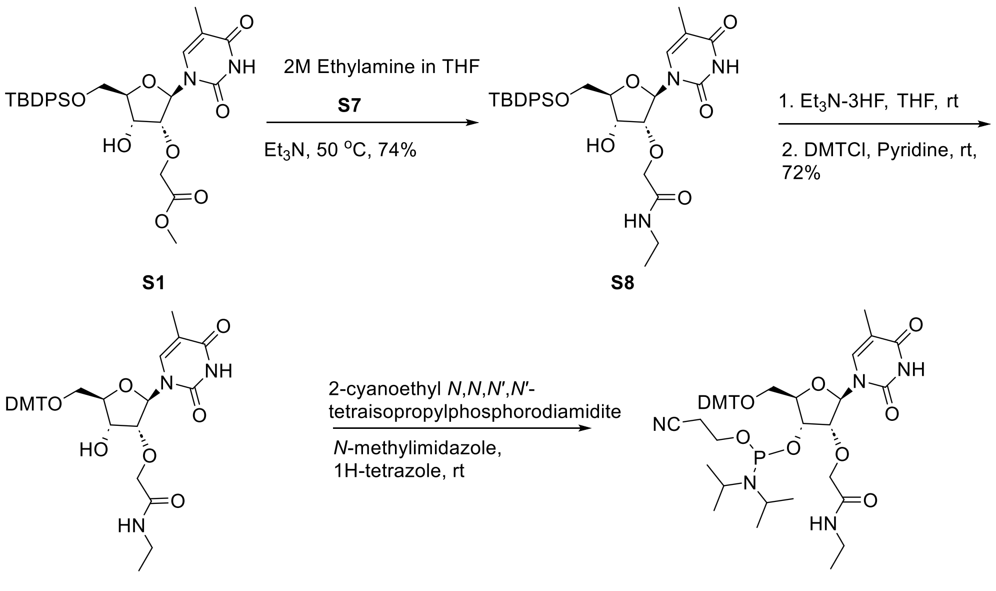

S1 S8 S9 S10

**Scheme S2**. Synthesis of 5'-*O*-DMT-2'-*O*-(NEA)-thymidine-3'-phosphoramidite **S10**

**Compound S9**. To a solution of compound **S8** (14.9 g, 25.6 mmol) in THF (250 mL) triethylamine (10.7 mL, 76.7 mmol) and triethylamine trihydrofluoride (8.3 mL, 51.2 mmol) were added. After stirring the reaction mixture for 12 h at room temperature and it was concentrated under reduced pressure. The residue was purified by silica gel column chromatography and eluted with 0-30% MeOH in the THF to yield a 5'-hydroxy analog of compound **S8 (**8.8 g). LRMS (ES, TOF): *m/z* calcd for $C_{14}H_{21}N_3NaO_7$: 366.1, found 366.1 $[M + 23]^+$. Dissolved 5'-hydroxy analog of compound **S8 (**8.8 g, 25.6 mmol) from above in pyridine (250 mL), and DMTCl (13.0 g, 38.4 mmol) was added. The reaction mixture was stirred at room temperature for 12 h and quenched with water (10 mL). Diluted the reaction mixture with ethyl acetate (150 mL) and washed with saturated aqueous sodium bicarbonate (350 mL), brine (350), dried ($Na_2SO_4$), filtered, and concentrated under reduced pressure. The crude product was purified by silica gel column chromatography and eluted with 1-8% MeOH in dichloromethane to yield **S9** (12.0 g, 72.4%).$^{1}$H NMR (300 MHz, $CDCl_3$): δ 10.45 (s, 1H), 7.76 (d, *J* = 1.2 Hz, 1H), 7.37-7.45 (m, 2H), 7.16-7.35 (m, 7H), 7.05 (t, *J* = 5.4 Hz, 1H), 6.72-6.93 (m, 4H), 5.87 (d, *J*= 1.5 Hz, 1H), 4.80 (d, *J* = 5.4 Hz, 1H), 4.29-4.56 (m, 3H), 4.15-4.20 (m, 1H), 3.95 (dd, *J* = 1.4, 5.4 Hz, 1H), 3.78 (d, *J* = 1.2 Hz, 6H), 3.44-3.61 (m, 2H), 3.15-3.42 (m, 2H), 1.35 (s, 3H), 1.13 (t, *J* = 7.23 Hz, 3H); $^{13}$C NMR (75 MHz, $CDCl_3$): δ 169.9, 164.0, 158.7, 151.3, 144.4, 135.4, 135.3, 135.1, 130.1, 128.1, 128.0, 127.1, 113.3, 111.1, 89.2, 86.7, 84.6, 83.7, 69.8, 68.0, 61.4, 60.4, 55.2, 34.2, 14.4, 11.9; LRMS (ESI TOF): *m/z* calcd for $C_{35}H_{39}N_3NaO_9$: 668.3, found 668.3 $[M + Na]^+$.

**5'-*O*-DMT-2'-*O*-(NEA)-thymidine-3'-phosphoramidite S10**. Compound **S9** (7.0 g, 10.8 mmol) was dissolved in anhydrous DMF (40 mL) and tetrazole (0.6 g, 8.7 mmol), *N*-methylimidazole (0.2 mL, 2.7 mmol) and 2-Cyanoethyl *N*,*N*,*N′*,*N′*-tetraisopropylphosphorodiamidite (5.2 mL, 16.3 mmol) were added at 0 $^{o}$C (ice bath). After removing the ice bath, the reaction was stirred for 12 hat rom temperature and diluted the reaction mixture with ethyl acetate (60 mL). The ethyl acetate solution was washed with saturated aqueous sodium bicarbonate solution (100 mL), brine (100 mL), dried ($Na_2SO_4$), and concentrated under reduced pressure. The residue was purified by silica gel column chromatography and eluted with 50-100% acetone in dichloromethane. Fractions were pooled together and concentrated under reduced pressure. The residue was dissolved in dichloromethane (10 mL) and added dropwise to vigorously stirred Hexanes (1.6 L). The precipitate formed was filtered and dried under reduced pressure to **yield S10** (9.2 g, <95%). $^{31}$P NMR (121MHz, $CDCl_3$): δ 151.0, 150.4; HRMS (ESI TOF): m/z calcd for $C_{44}H_{56}N_5O_{10}P$: 845.3765, found 644.3678 $[M - H]^-$.

**Compound S11**. To a solution of compound **S1** (21.4 g, 36.7 mmol) in methanol (80 mL), propylamine (30.8 mL, 367.0 mmol) was added, and the mixture was stirred at 80 $^{o}$C for two days. The reaction mixture was concentrated at reduced pressure. The residue was purified by silica gel column chromatography and eluted first with 20-30% ethyl acetate in dichloromethane (4-5 column volume) and then with 0-5% MeOH in dichloromethane to yield **S11** (11.3 g, 52%). $^{1}$H NMR (300 MHz, $CDCl_3$): δ 10.15 (br s, 1H), 7.61-7.77 (m, 4H), 7.60 (s, 1H), 7.33-7.50 (m, 6H), 6.99 (t, $J$ = 5.6 Hz, 1H), 5.94 (d, $J$ = 2.8 Hz, 1H), 4.25-4.54 (m, 4H), 4.08-4.22 (m, 2H), 3.79-4.00 (m, 2H), 3.07-3.34 (m, 2H), 1.43-1.62 (m, 5H), 1.05-1.18 (m, 9H), 0.83-0.97 (m, 3H); LRMS (ESI TOF): *m/z* calcd for $C_{31}H_{41}N_3NaO_7Si$: 618.3 , found 618.3 $[M + Na]^+$.

**Compound S12**. Compound **S11** (11.3 g, 19.0 mmol) was dissolved in THF (130 mL), triethylamine (8.3 mL, 59.4 mmol), and triethylamine trihydrofluoride (8.4 mL, 51.5 mmol) was added. The mixture was stirred at room temperature for 12 h and concentrated under reduced pressure. The residue was purified by silica gel column chromatography and eluted with 0-10% MeOH in THF to give **S12** (6.8 g, 95%). $^{1}$H NMR (300 MHz, DMSO-$d_6$): δ11.33 (s, 1H), 7.88 (t, $J$ = 5.8 Hz, 1H), 7.80 (d, $J$ = 1.2 Hz, 1H), 5.83 (d, $J$ = 4.0 Hz, 1H), 5.48 (d, $J$ = 6.1 Hz, 1H), 5.18 (t, $J$ = 5.1 Hz, 1H), 4.07-4.16 (m, 1H), 4.03-4.16 (m, 2H), 3.93-4.00 (m, 1H), 3.84-3.92 (m, 1H), 3.66-3.79 (m, 1H), 3.54-3.64 (m, 1H), 2.95-3.13 (m, 2H), 1.76 (d, $J$ = 1.0 Hz, 3H), 1.36-1.52 (m, 2H), 0.84 (t, $J$ = 7.4 Hz, 3H); LRMS (ESI TOF): *m/z* calcd for $C_{15}H_{24}N_3O_7$: 357.2, found 358.2 $[M + H]^+$.

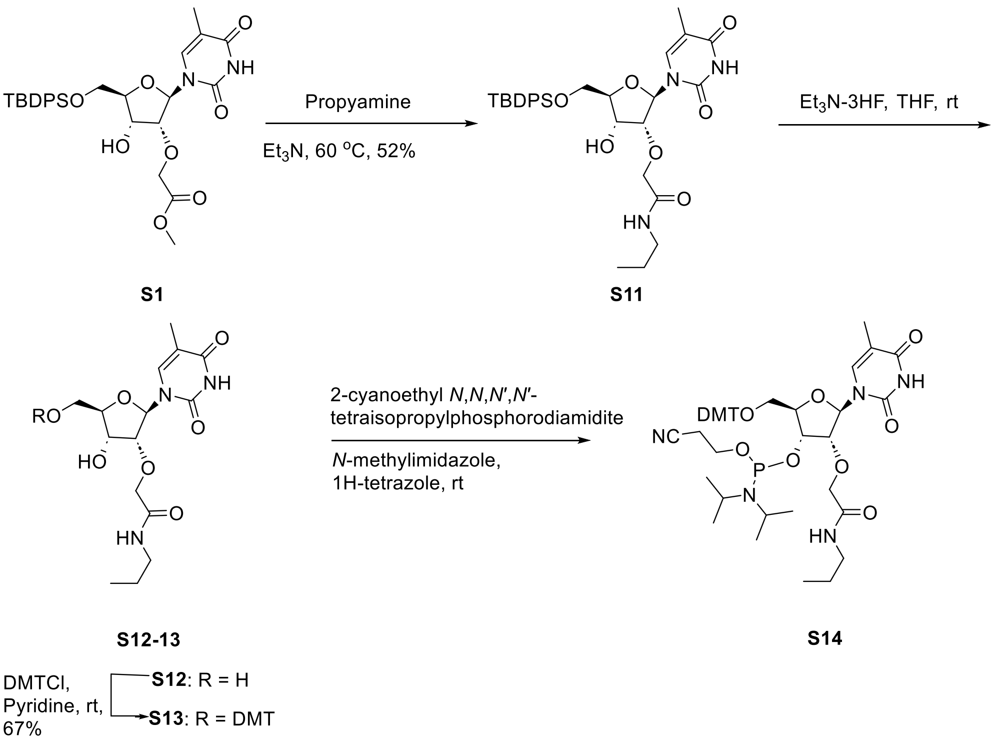

**Scheme S3**. Synthesis of 5’-*O*-DMT-2’-*O*-(NPA)-thymidine-3’-phosphoramidite **S14**

**Compound S13**. To a solution of **S12** (6.9 g, 19.3 mmol) in anhydrous pyridine (200 mL) DMTCl (9.8 g, 29.0 mmol) was added at room temperature. The reaction mixture was stirred at room temperature for 12 h and quenched with water (20 mL). The reaction mixture was diluted ethyl with ethyl acetate (200 mL) and washed with saturated aqueous sodium bicarbonate (400 mL), brine (400), dried ($Na_2SO_4$), filtered, and concentrated under reduced pressure. The residue was purified by silica gel column chromatography and eluted first with 0-50% ethyl acetate in a hexanes (4 column volume) and then with 0-5% MeOH in dichloromethane to yield **S13** (8.5 g, 67%).$^{1}$H NMR (300 MHz, $CDCl_3$): δ 10.36 (s, 1H), 7.74 (s, 1H), 7.37-7.46 (m, 2H), 7.18-7.36 (m, 7H), 7.03 (t, *J* = 5.7 Hz, 1H), 6.83 (d, *J* = 6.0 Hz, 4H), 5.87 (d, *J* = 1.5 Hz, 1H), 4.71 (d, *J* = 5.1 Hz, 1H), 4.30-4.54 (m, 3H), 4.19 (d, *J* = 7.8 Hz, 1H), 3.95 (dd, *J* = 1.5, 5.4 Hz, 1H), 3.78 (s, 6H), 3.43-3.62 (m, 2H), 3.03-3.37 (m, 2H), 1.44-1.61 (m, 2H), 1.37 (s, 3H), 0.91 (t, *J* = 7.4 Hz, 3H);

$^{13}C$ NMR (75 MHz, $CDCl_3$): δ 170.0, 164.0, 158.7, 151.3, 144.5, 135.4, 135.3, 135.1, 130.1, 128.1, 128.0, 127.0, 113.3, 111.1, 89.2, 86.7, 84.6, 83.7, 69.8, 68.1, 61.5, 55.2, 41.1, 22.6, 11.9, 11.4; LRMS (ESI TOF): *m/z* calcd for $C_{36}H_{41}N_3NaO_9$: 682.3, found 682.3 $[M + 23]^+$.

**5'-*O*-DMT-2'-*O*-(NPA)-thymidine-3'-phosphoramidite S14.** Compound **S13** (5.9 g, 8.9 mmol) was dissolved in anhydrous DMF (43.6 mL). To this tetrazole (0.5 g, 7.1 mmol), N-methyl imidazole (0.2 mL, 2.2 mmol) and 2-Cyanoethyl *N*,*N*,*N′*,*N′*-tetraisopropylphosphorodiamidite (4.2 mL, 13.3 mmol) were added when it was placed in an ice bath. The reaction mixture was removed from the ice bath and stirred for 2 h at room temperature. Diluted the reaction mixture with ethyl acetate (60 mL) and washed by saturated aqueous sodium bicarbonate solution (150 mL), brine (150 mL), dried ($Na_2SO_4$) and concentrated under reduced pressure. The crude product was purified by silica gel column chromatography and eluted with 20-50% acetone in a dichloromethane. Fractions were pooled together and concentrated under reduced pressure. The residue was dissolved in dichloromethane (10 mL) and added dropwise to vigorously stirred hexanes (1 L). The precipitate formed was collected by filtration and dried under reduced pressure to yield Compound **S14** (7.6 g, <95%). $^{31}P$ NMR (121MHz, $CDCl_3$): δ 151.0, 150.5; HRMS (ESI, TOF): *m/z* calcd for $C_{45}H_{57}N_5O_{10}P$: 859.3921, found 858.3839 $[M - H]^-$.

**Compound S16.** Compound **S1** (20.4 g, 35.1 mmol), dissolved in methanol (100 mL). To this cyclopropylamine **S15** (24.3 mL, 351.0 mmol) was added and the reaction mixture was stirred 80 $^oC$ for 12 h. The reaction mixture was concentrated under reduced pressure. The residue was purified by silica gel column chromatography and eluted first with 0-50% ethyl acetate in hexanes (4-5 column volume) and then with 0-5% MeOH in dichloromethane to yield **16** (14.8 g, 71%). $^1H$ NMR (300 MHz, $CDCl_3$): δ 10.43 (s, 1H), 7.62-7.72 (m, 5H), 7.61 (s, 1H), 7.33-7.50 (m, 6H), 7.22 (d, *J* = 3.7 Hz, 1H), 5.90 (d, *J* = 2.4 Hz, 1H), 4.68 (d, *J* = 5.1 Hz, 1H), 4.23-4.50 (m, 3H),

4.04-4.22 (m, 2H), 3.84-4.02 (m, 2H), 2.62-2.84 (m, 1H), 1.48 (s, 3H), 1.03-1.22 (m, 9H), 0.65-0.86 (m, 2H), 0.41-0.61 (m, 2H); $^{13}$C NMR (75 MHz, $CDCl_3$): δ 171.4, 163.9, 151.4, 135.5, 135.2, 134.8, 133.2, 132.4, 130.1, 130.0, 128.0, 127.9, 111.3, 88.8, 84.6, 84.6, 69.8, 67.6, 62.3, 62.1, 22.3, 27.0, 19.5, 11.9, 6.3, 6.1, 5.9; LRMS (ESI TOF): *m/z* calcd for $C_{31}H_{39}N_3NaO_7Si$: 593.3, found 616.3 $[M + Na]^+$.

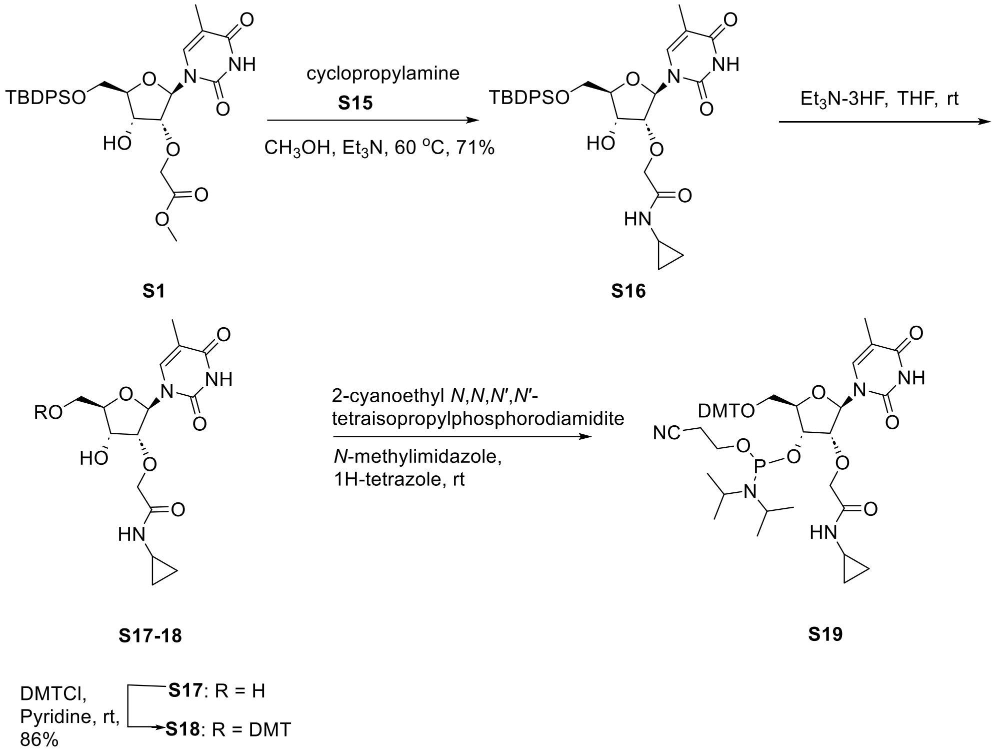

**Scheme S4**. Synthesis of 5'-*O*-DMT-2'-*O*-(NcPA)-thymidine-3'-phosphoramidite **S19**

**Compound 17**. To a THF solution (250 mL) of **S16** (14.8 g, 24.9 mmol), triethylamine (13.9 mL, 99.5 mmol), triethylamine trihydrofluoride (12.2 mL, 74.6 mmol) was added. After stirring for three days at room temperature, the reaction mixture was concentrated under reduced pressure. The residue after purification by silica gel column chromatography (5-10% MeOH in the THF/dichloromethane (1:1) as eluent) yield **S17** (8.8 g, <95%). $^1$H NMR (300 MHz, DMSO-$d_6$): δ 11.18 (s, 1H), 7.76-7.90 (m, 1H), 7.75 (s, 1H), 5.82 (d, $J$ = 4.1 Hz, 1H), 5.38 (d, $J$ = 5.9 Hz, 1H),

5.02 (t, *J* = 4.9 Hz, 1H), 3.84-4.17 (m, 5H), 3.56-3.80 (m, 2H), 2.60-2.76 (m, 1H), 1.77 (s, 3H), 0.60-0.72 (m, 2H), 0.37-0.51 (m, 2H); $^{13}$C NMR (75 MHz, DMSO-d$_6$): δ 169.9, 163.4, 150.3, 135.7, 108.9, 86.5, 84.2, 82.4, 69.2, 68.1, 60.0, 21.7, 11.9, 5.3; LRMS (ESI TOF): *m/z* calcd for $C_{15}H_{21}N_3NaO_7$: 378.1 , found 378.1 $[M + Na]^+$.

**Compound 18**. Compound **S17** (3.9 g, 10.9 mmol) was dissolved in anhydrous pyridine (110 mL). To this DMTCl (4.4 g, 13.1 mmol) was added and the reaction was stirred at room temperature for 12 h. After quenching the reaction with water (10 mL) the mixture was diluted with ethyl acetate (200 mL). It was then washed with aqueous sodium bicarbonate solution (300 mL), brine (300 mL), and dried ($Na_2SO_4$), filtered, and concentrated under reduced pressure. The residue c was purified by silica gel column chromatography (eluted with 0-5% MeOH in a dichloromethane) to yield **S18** (6.2 g, 86%). $^1$H NMR (300 MHz, $CDCl_3$): δ10.52 (s, 1H), 7.77 (d, *J* = 1.2 Hz, 1H), 7.36-7.51 (m, 2H), 7.18-7.35 (m, 7H), 6.83 (dd, *J* = 0.83, 8.90 Hz, 4H), 5.84 (d, *J* = 1.3 Hz, 1H), 4.77 (d, *J* = 5.4 Hz, 1H), 4.27-4.57 (m, 3H), 4.19 (d, *J* = 8.1 Hz, 1H), 3.92 (dd, *J* = 1.1, 5.4 Hz, 1H), 3.78 (d, *J* = 1.2 Hz, 6H), 3.38-3.64 (m, 2H), 2.61-2.85 (m, 1H), 1.37 (d, *J* = 0.8 Hz, 3H), 0.64-0.85 (m, 2H), 0.37-0.60 (m, 2H); $^{13}$C NMR (75 MHz, $CDCl_3$): δ 171.6, 164.1, 158.7, 151.4, 144.5, 135.5, 135.3, 135.2, 130.1, 128.1, 128.0, 127.1, 113.3, 111.1, 89.4, 86.7, 84.7, 83.7, 69.7, 67.8, 61.4, 55.2, 22.4, 11.9, 6.2, 5.9; LRMS (ESI TOF): *m/z* calcd for $C_{36}H_{39}N_3NaO_9$: 680.3, found 680.2 $[M + Na]^+$.

**5'-*O*-DMT-2'-*O*-(NcPA)-thymidine-3'-phosphoramidite S19.** To a DMF (48 mL) solution of compound **S18** (6.5 g, 9.8 mmol), tetrazole (0.6 g, 7.9 mmol), *N*-Methylimidazole (0.24 mL, 2.5 mmol) and 2-cyanoethyl *N*,*N*,*N′*,*N′*-tetraisopropylphosphorodiamidite (4.7 mL, 14.7 mmol) was added. After stirring for two h at room temperature, the reaction mixture was diluted with ethyl acetate (100 mL) and washed with saturated aqueous sodium bicarbonate solution (200 mL), brine (200 mL), dried ($Na_2SO_4$), filtered and concentrated at reduced pressure. Fraction obtained from the purification of the residue by silica gel column chromatography (20-50% acetone in a dichloromethane as eluent) were concentrated. The residue was dissolved in dichloromethane (10 mL) and added dropwise to vigorously stirred hexanes (1.5 L). The precipitate formed was collected by filtration and dried under reduced pressure to give **S19** (7.9 g, 93.5%). $^{31}$P NMR (121 MHz, $CDCl_3$): δ 151.0, 150.4; HRMS (ESI TOF): *m/z* calcd for $C_{45}H_{55}N_5O_{10}P$: 857.3765, found 856.3680 $[M - H]^-$.

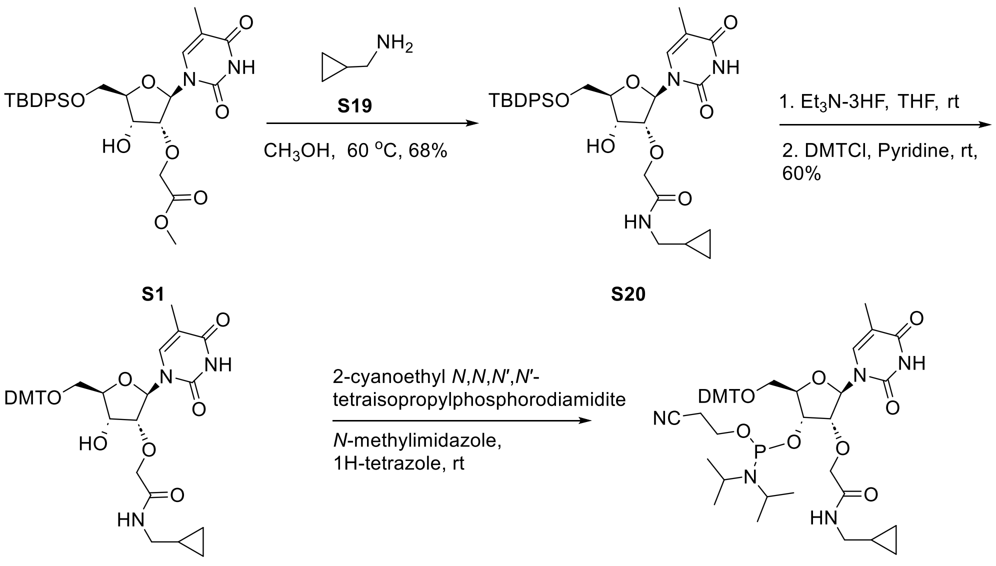

**Scheme S5**. Synthesis of 5'-*O*-DMT-2'-*O*-(McPA)-thymidine-3'-phosphoramidite **S22**

**Compound S20**. Compound **S1** (20.6 g, 35.3 mmol) was dissolved in methanol solution (150 mL). To this cyclopropane methylamine hydrochloride **S19** (40.0 g, 368.0 mmol) was added and stirred at 60 °C for three days. The reaction mixture was concentrated under reduced pressure and the residue was purified by silica gel column chromatography (elute first with 30% ethyl acetate in hexanes (4-5 column volume) and then with 5% MeOH in dichloromethane) to yield **S20** (14.6 g, 68%). $^1$H NMR (300 MHz, $CDCl_3$): δ 7.62-7.74 (m, 4H), 7.59 (d, $J$ = 1.2 Hz, 1H), 7.31-7.50 (m,

6H), 7.18 (t, *J* = 5.4 Hz, 1H), 6.76 (br s, 1H), 5.96 (d, *J* = 2.9 Hz, 1H), 4.27-4.50 (m, 3H), 4.08-4.24 (m, 2H), 3.94 (dt, *J* = 2.4, 5.9 Hz, 2H), 2.98-3.26 (m, 3H), 1.49 (d, *J* = 1.0 Hz, 3H), 1.09 (s, 9H), 0.82-1.03 (m, 1H), 0.37-0.60 (m, 2H), 0.08-0.30 (m, 2H); $^{13}$C NMR (75 MHz, $CDCl_3$): δ 171.7, 169.7, 163.9, 151.3, 135.5, 135.2, 134.7, 133.1, 132.3, 130.1, 130.0, 128.0, 127.9, 111.3, 88.3, 84.6, 84.4, 69.9, 68.0, 62.5, 62.1, 44.1, 43.8, 27.0, 19.4, 11.9, 10.6, 10.5, 3.5, 3.4, 3.4; LRMS (ESI TOF): *m/z* calcd for $C_{32}H_{41}N_3NaO_7Si$: 630.3, found 630.3 $[M + Na]^+$.

**Compound S21**. To a solution of **S20** (15.3 g, 25.1 mmol) in THF (200 mL) triethylamine (13.5 mL, 96.8 mmol)) and triethylamine trihydrofluoride (13.5 mL, 96.8 mmol) was added. After stirring for 12 h at room temperature, the reaction mixture was concentrated under reduced pressure. The residue obtained was purified by silica gel column chromatography (eluent 0-30% MeOH in the dichloromethane) to yield 5'-hydroxyl analog of **S20** (9.38 g). LRMS (ESI TOF): m/z calcd for $C_{16}H_{23}N_3NaO_7$: 392.2, found 392.1 $[M + Na]^+$. The 5'-hydroxyl analog of **S20** (9.38 g, 25.1 mmol) from above was dissolved in anhydrous pyridine (250 mL), DMTCl (17.1 g, 50.4 mmol) was added, and the reaction was stirred at room temperature for 12 h. The reaction was quenched by adding methanol (5 mL) diluted with ethyl acetate (300 mL) and washed with saturated aqueous sodium bicarbonate (300 mL), brine (300 mL), and dried ($Na_2SO_4$) filtered and concentrated under reduced pressure. The crude product was purified by silica gel column chromatography and eluted first with 8-50% ethyl acetate in a dichloromethane (4 column volume) and then 5-8% MeOH in a dichloromethane to yield **S21** (10.2 g, 60%).$^1$H NMR (300 MHz, $CDCl_3$): δ 10.35 (s, 1H), 7.74 (d, *J* = 1.2 Hz, 1H), 7.36-7.48 (m, 2H), 7.18-7.36 (m, 7H), 7.03-7.16 (m, 1H), 6.83 (dd, *J* = 0.9, 8.96 Hz, 4H), 5.88 (d, *J* = 1.7 Hz, 1H), 4.70 (d, *J* = 5.5 Hz, 1H), 4.31-4.55 (m, 3H), 4.19 (d, *J* = 7.6 Hz, 1H), 3.97 (dd, *J* = 1.7, 5.4 Hz, 1H), 3.78 (s, 6H), 3.43-3.62 (m, 2H), 2.95-3.25 (m, 2H), 1.37 (d, *J* = 0.9 Hz, 3H), 0.79-1.03 (m, 1H), 0.38-0.54 (m, 2H),

0.14-0.27 (m, 2H); $^{13}C$ NMR (75 MHz, $CDCl_3$): δ 169.9, 164.0, 158.7, 158.7, 151.3, 144.4, 135.5, 135.3, 135.1, 130.1, 130.1, 128.1, 128.0, 127.1, 113.3, 111.1, 89.1, 86.7, 84.6, 83.6, 69.9, 68.1, 61.5, 55.2, 44.2, 11.9, 10.5, 3.5, 3.5; LRMS (ESI TOF): *m/z* calcd for $C_{37}H_{41}N_3NaO_9$: 694.3, found 694.3 $[M + Na]^+$.

**5'-*O*-DMT-2'-*O*-(McPA)-thymidine-3'-phosphoramidite S22.** To a DMF (50 mL) solution of compound **S21** (7.1 g, 10.6 mmol) tetrazole (0.6 g, 8.4 mmol)) *N*-methylimidazole (0.2 mL, 2.6 mmol) 2-cyanoethyl *N*,*N*,*N′*,*N′*-tetraisopropylphosphorodiamidite (4.8 mL, 15.8 mmol) at 0 °C (ice bath). The reaction mixture was removed from the ice bath and stirred at room temperature for 12 h. Diluted the reaction mixture with ethyl acetate (100 mL) and washed with saturated aqueous sodium bicarbonate solution (100 mL), brine (100 mL), dried ($Na_2SO_4$) filtered and concentrated under reduced pressure. The crude product was purified by silica gel column chromatography (50-70% ethyl acetate in a hexanes). Fraction-containing desired products (assessed by LC-MS analysis) were pooled together and concentrated under reduced pressure, and the residue was dissolved in dichloromethane (10 mL) and dropwise added to vigorously stirred hexanes (2 L). The precipitate formed was collected by filtration and dried under reduced pressure to yield **S22** (9.1 g, 99%). $^{31}P$ NMR (121MHz, $CDCl_3$): δ 150.9, 150.5; HRMS (ESI TOF): *m/z* calcd for $C_{46}H_{57}N_5O_{10}P$: 871.3921, found 870.3839 $[M - H]^-$.

**Compound S23**. The compound **S5** (3.5 g, 5.4 mmol) was dissolved in anhydrous DMF (36 mL). Imidazole (2.2 g, 32.5 mmol) and tert-butyl-chlorodimethylsilane (3.3 g, 21.7 mmol) were added and stirred at room temperature for 12 h. MeOH (3 mL) was added and stirred for 30 min at room temperature to quench the reaction. The reaction mixture was diluted with ethyl acetate (60 mL) and washed with aqueous sodium bicarbonate (100 mL), brine (100 mL), and dried ($Na_2SO_4$) filtered and concentrated under reduced pressure. The crude product was purified by silica gel column chromatography (0-5% methanol in a dichloromethane) to yield **S23** (4.06 g, 98.6%). LRMS (ESI TOF): *m/z* calcd for $C_{41}H_{53}N_3NaO_9Si$: 782.3, found 782.3 $[M + Na]^+$.

**Compound S24:** To a suspension of 1,2,4-triazole (9.4 g, 136.0 mmol) in anhydrous acetonitrile (70 mL) with an ice-bath, $POCl_3$ (3.1 mL, 33.9 mmol) was slowly added. The mixture was stirred at 0 $^oC$ for 15 minutes, and triethylamine (23.6 mL, 169.0 mmol) was introduced. After stirring the reaction mixture for 30 min at 0 $^oC$ (ice-bath) a solution of **S23** (3.2 g, 4.2 mmol) in acetonitrile (20 mL) was added. The reaction mixture was removed from the ice-bath and stirred atfor 12 h at room temperature. The reaction mixture was diluted with ethyl acetate (200 mL) and washed with water (250 mL), dried ($Na_2SO_4$), filtered, and concentrated to give a triazolide analog of **S23**. LRMS (ESI TOF): *m/z* calcd for $C_{43}H_{54}N_6NaO_8Si$: 833.4, found 833.3 $[M + Na]^+$. To a 1,4 dioxane (31 mL) solution of triazolide analog of **S23** (3.4 g, 4.2 mmol), aqueous ammonia (17.2 mL, 28-30 wt%) was added and stirred at room temperature for two h in a sealed bottle. The reaction mixture was concentrated under reduced pressure and dried over $P_2O_5$ under reduced pressure overnight to yield a 5-methylcytidine analog of compound **S23** (3.2 g). LRMS (ESI TOF): *m/z* calcd for $C_{41}H_{54}N_4NaO_8Si$: 781.3, found 781.3 $[M + Na]^+$. The cytidine analog of compound **S23** (3.2 g, 4.2 mmol) from above was dissolved in anhydrous DMF (42 mL) and isobutyric anhydride (1.1 mL, 7.64 mmol) was added. After stirring for 18 h, the reaction was quenched by adding MeOH (4 mL) and stirring continued for 10 min. Dilute the reaction mixture with ethyl acetate (60 mL), wash it with brine (100 mL), dry it ($Na_2SO_4$), filter it, and concentrate it under reduced pressure. The crude product was purified by silica gel column chromatography and eluted with 0-50% acetone in dichloromethane to yield **S24** (2.8 g, 81%). $^1H$ NMR (300 MHz, DMSO-$d_6$): δ 9.97 (br s, 1H), 7.99 (br s, 1H), 7.15-7.48 (m, 9H), 6.89 (d, $J$ = 8.2 Hz, 4H), 5.92 (d, $J$ = 2.1 Hz, 1H), 4.22-4.50 (m, 4H), 4.12-4.25 (m, 1H), 4.05 (d, $J$ = 5.9 Hz, 1H), 3.74 (s, 6H), 3.09-3.54 (m,

3H), 2.89 (s, 1H), 2.78 (s, 3H), 1.49 (s, 3H), 1.09 (s, 3H), 1.07 (s, 3H), 0.74 (s, 9H), 0.02-0.12 (m, 3H), -0.05 (s, 3H); $^{13}$C NMR (75 MHz, DMSO-d$_6$): δ 168.0, 162.3, 158.2, 144.3, 135.1, 135.0, 129.8, 129.8, 127.9, 127.8, 126.9, 113.2, 86.0, 68.6, 55.0, 54.9, 35.8, 35.7, 34.8, 30.7, 25.6, 25.5, 19.5, 19.2, 19.0, 17.7, 17.6, -4.6, -5.3; LRMS (ESI TOF): *m/z* calcd for $C_{45}H_{60}N_4NaO_9Si$: 851.4, found 851.4 $[M + Na]^+$.

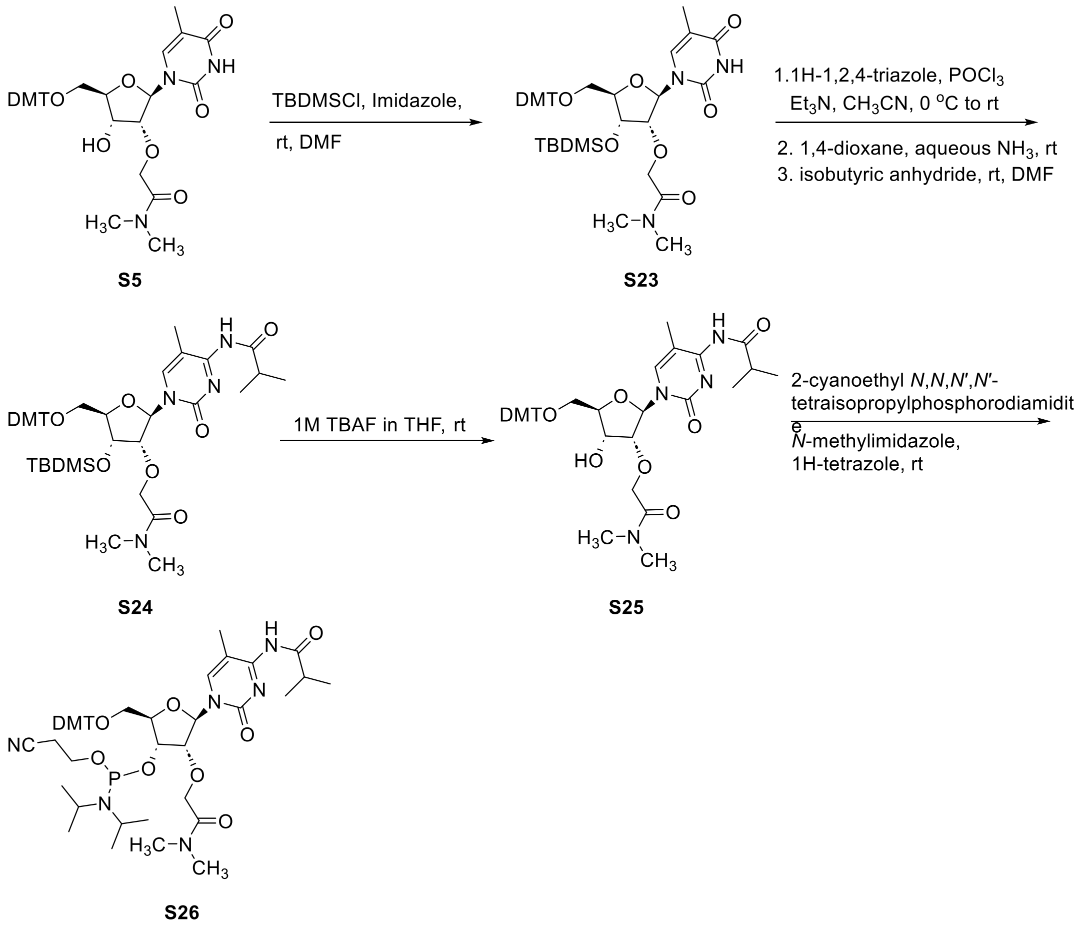

**Scheme S6**. Synthesis of 5’-*O*-DMT-2’-*O*-(DMA)-*N*-4-isobutyryl-5-methyl-cytidine-3’-phosphoramidite **S26**

**Compound 25.** To a THF solution (34 mL) of compound **S24** (2.8 g, 3.4 mmol), 1N TBAF in THF (5.1 mL,5.1 mmol) was added. The reaction mixture was stirred at room temperature for 3 h and concentrated under reduced pressure. The residue was re-dissolved in ethyl acetate (60 mL), washed with brine (100 mL), dried (Na2SO4), filtered, and concentrated under reduced pressure. The crude product was purified by silica gel column chromatography (0-50% acetone in dichloromethane) to yield **S25** (2.3 g, 94%). $^{1}$H NMR (300 MHz, DMSO-$d_6$): δ 10.04 (br s, 1H), 7.86 (s, 1H), 7.19-7.44 (m, 9H), 6.91 (d, $J$ = 9.0 Hz, 4H), 5.85-5.95 (m, 2H), 4.40-4.56 (m, 2H), 4.16-4.20 (m, 1H), 4.01-4.16 (m, 2H), 3.68-3.84 (m, 6H), 3.25-3.40 (m, 3H), 2.90 (s, 3H), 2.83 (s, 3H), 1.50 (s, 3H), 1.09 (s, 3H), 1.07 (s, 3H); $^{13}$C NMR (75 MHz, DMSO-$d_6$): δ 169.5, 158.2, 144.6, 144.5, 135.4, 135.1, 129.7, 129.7, 128.0, 127.7, 126.8, 113.3, 86.0, 85.8, 83.3, 83.2, 68.6, 68.5, 68.0, 62.4, 55.0, 54.9, 53.2, 35.4, 35.0, 29.6, 28.9, 20.0, 19.2, 19.1, 13.9, 13.5, 11.6; LRMS (ESI TOF): *m/z* calcd for $C_{39}H_{47}N_4O_9$: 714.3, found 715.3 $[M + H]^+$.

**5'-*O*-DMT-2'-*O*-(DMA)-*N*-4-isobutyryl-5-methyl-cytidine-3'-phosphoramidite S26.** Compound **S25** (2.1 g, 2.9 mmol) and diisopropyl ammonium tetrazolide (0.5 g, 2.8 mmol) were placed in a flask with a stir bar and under reduced pressure over $P_2O_5$ for three days. The mixtures were suspended in anhydrous acetonitrile (19 mL). To this suspension, 2-cyanoethyl *N*,*N*,*N′*,*N′*-tetraisopropylphosphorodiamidite (1.4 mL, 4.3 mmol) was added. The reaction mixture was stirred at room temperature for 4 h and filtered. The filtrate was concentrated under reduced pressure. The crude product was purified by silica gel column (primed with 1% triethylamine in dichloromethane) and eluted with 5-20% acetone in dichloromethane containing 1% triethylamine. The fraction containing desired products were pooled together and concentrated under reduced pressure. The residue was dissolved in dichloromethane (10 mL) and dropped into hexanes (1.5 L) with vigorous stirring. The precipitate formed was collected by filtration to yield **S26** (2.3 g, 85%). $^{31}$P NMR (121 MHz, DMSO-$d_6$) δ: 149.75, 149.62; HRMS (ESI TOF): *m/z* calcd for $C_{48}H_{62}N_6O_{10}P$: 914.4343, found 913.4263 $[M - 1]^-$.

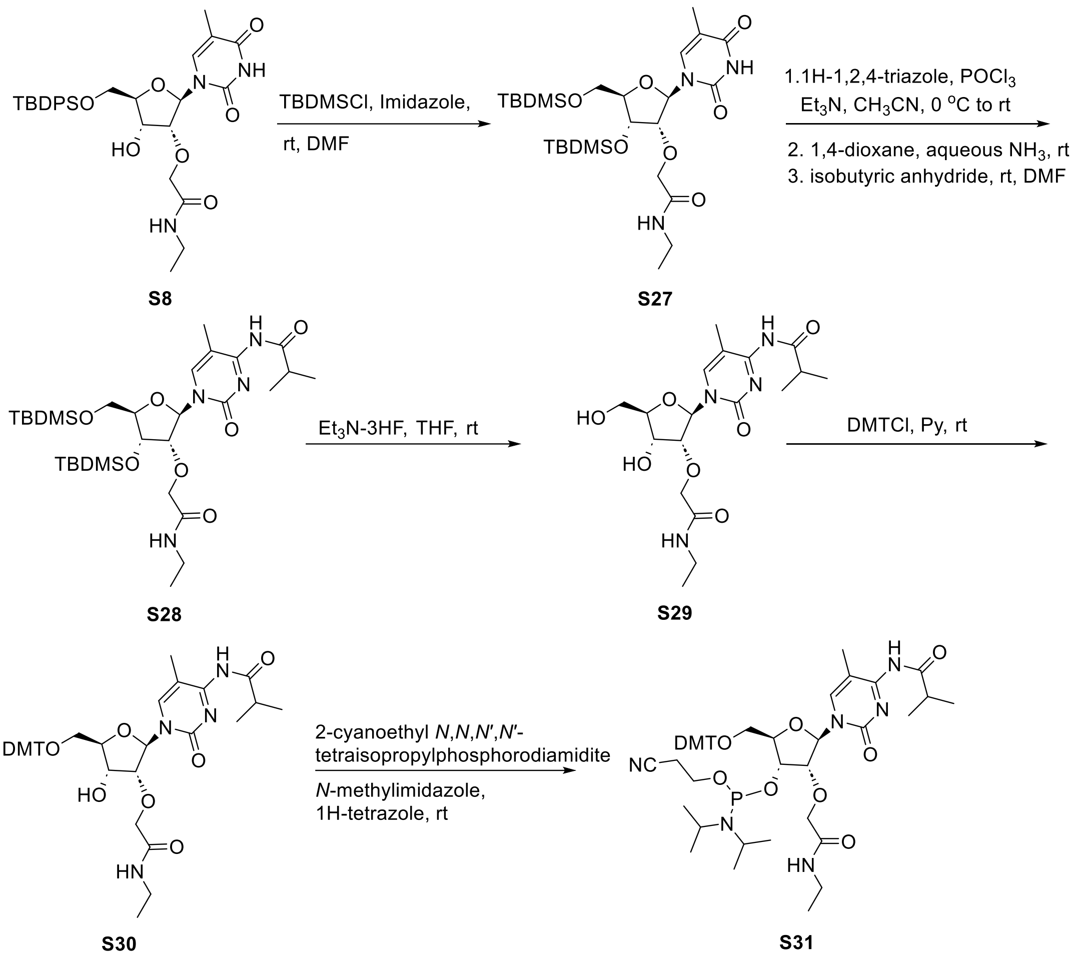

**Scheme S7**. Synthesis of 5'-*O*-DMT-2'-*O*-(NEA)-*N*-4-isobutyryl-5-methyl-cytidine-3'-phosphoramidite **S31**

**Compound S27**. To a DMF solution (122 mL) of compound **S8** (10.9 g, 18.7 mmol) imidazole (7.6 g, 112.0 mmol), *tert*-butyl-chlorodimethylsilane (11.3 g, 74.9 mmol) were added. After stirring the reaction mixture for 12 h at room temperature, the reaction was quenched by adding MeOH (3 mL) and stirring continued for 30 min. The reaction mixture was diluted with ethyl acetate (150 mL) washed with brine (200 mL), dried ($Na_2SO_4$), filtered, and concentrated under reduced pressure. The crude product was purified by silica gel column chromatography (0-100% ethyl acetate in hexanes) to yield **S27** (9.4 g, 72%). $^1$H NMR (300 MHz, DMSO-d$_6$): δ 11.42 (br s, 1H), 7.64 (dt, $J$ = 1.6, 8.1 Hz, 4H), 7.30-7.56 (m, 8H), 5.91 (d, $J$ = 4.9 Hz, 1H), 4.38 (t, $J$ = 4.9

Hz, 1H), 4.18 (t, $J$ = 4.9 Hz, 1H), 3.88-4.08 (m, 4H), 3.70-3.81 (m, 1H), 2.99-3.18 (m, 2H), 1.49 (d, $J$ = 0.9 Hz, 3H), 0.95-1.05 (m, 12H), 0.84 (s, 9H), 0.07 (s, 3H), 0.05 (s, 3H); $^{13}$C NMR (75 MHz, DMSO-d$_6$): δ 168.0, 163.7, 150.5, 135.8, 135.0, 132.5, 132.3, 130.1, 130.0, 128.0, 127.9, 109.5, 86.7, 83.9, 80.3, 69.7, 69.1, 63.1, 59.733.0, 26.7, 25.6, 20.7, 18.8, 17.7, 14.7, 14.1, 11.8, -4.8, -5.1; LRMS (ESI TOF): *m/z* calcd for $C_{36}H_{53}N_3NaO_7Si_2$: 718.3, found 718.3 $[M + Na]^+$.

**Compound S28**. To 1,2,4-triazole (29.4 g, 426.0 mmol) suspension in acetonitrile (130 mL) in an ice-bath, $POCl_3$ (9.7 mL, 107.0 mmol) was slowly added. The mixture was stirred at 0 $^o$C for 15 min and triethylamine (74.3 mL, 533.0 mmol) was introduced. The reaction was stirred at 0 $^o$C for 30 min and a solution of compound **S27** (9.3 g, 13.3 mmol) in acetonitrile (40 mL) was added. The reaction mixture was removed from the ice bath and stirred at room temperature for 18 h. The reaction was quenched with water (10 mL) and diluted with ethyl acetate (200 mL) and washed with water (300 mL), dried ($Na_2SO_4$), filtered, and concentrated under reduced pressure to yield a triazolide analog of compound **S27** (10.6 g). LRMS (ESI TOF): *m/z* calcd for $C_{38}H_{54}N_6NaO_6Si_2$: 769.3, found 769.3 $[M + Na]^+$. To a 1,4 dioxane (31 mL) solution of triazolide analog of compound **S27** (10.0 g, 13.3 mmol), aqueous ammonia (54.3 mL, 28-20 wt%) was added. The reaction vessel was sealed with a septum and stirred at room temperature for 12 h. The reaction mixture was concentrated under reduced pressure to yield 5-methylcytidine analog of **S28** (9.3 g). LRMS (ESI TOF): *m/z* calcd for $C_{36}H_{54}N_4NaO_7Si_2$: 717.3, found 717.3 $[M + Na]^+$. 5-Methylcytidine analog of **S27** (1.4 g, 1.9 mmol) from above was dried over $P_2O_5$ under reduced pressure overnight and dissolved in anhydrous DMF (19 mL) and isobutyric anhydride (0.4 mL, 2.9 mmol) was added. After stirring at room temperature for 5 h, the reaction was quenched by adding MeOH (0.5 mL) and continued stirring for 10 min. The reaction mixture was diluted with ethyl acetate (100 mL) and washed with brine (100 mL), dried ($Na_2SO_4$), filtered and concentrated under reduced pressure. The crude product was purified by silica gel column chromatography (0-100% ethyl acetate in a dichloromethane) to yield **S28** (1.1 g, 74%). $^1$H NMR (300 MHz, DMSO-$d_6$): δ 10.03 (br s, 1H), 7.78 (br s, 1H), 7.66 (m, 4H), 7.56 (t, *J* = 5.8 Hz, 1H), 7.36-7.52 (m, 6H), 5.91 (d, *J* = 2.7 Hz, 1H), 4.35 (dd, *J* = 5.3, 6.8 Hz, 1H), 3.80-4.21 (m, 6H), 2.98-3.22 (m, 2H), 2.80-2.98 (br s, 1H), 1.54 (s, 3H), 0.97-1.11 (m, 18H), 0.83 (s, 9H), 0.06 (s, 3H), 0.03 (s, 3H); $^{13}$C NMR (75 MHz, DMSO-$d_6$): δ 168.1, 135.0, 134.9, 132.7, 132.4, 130.1, 130.0, 128.0, 127.9, 89.0, 83.4, 81.5, 69.4, 69.2, 62.6, 55.8, 33.0, 29.6, 26.8, 25.5, 19.1, 19.1, 19.0, 17.6, 14.7, -4.7, -5.2; LRMS (ESI TOF): *m/z* calcd for $C_{40}H_{60}N_4NaO_7Si_2$: 787.4, found 787.4 $[M + Na]^+$.

**Compound S29**. To a THF solution (23 mL) of compound **S28** (2.1 g, 2.7 mmol) triethylamine (1.3 mL, 9.3 mmol)) and triethylamine trihydrofluoride (1.3 mL, 8.1 mmol) was added. The mixture was stirred at room temperature for 12 h and concentrated under reduced pressure. The residue was purified by silica gel column chromatography (0-20% MeOH in the dichloromethane) to yield **S29** (1.1 g, <95%). $^1$H NMR (300 MHz, DMSO-$d_6$): δ 10.25 (br s, 1H), 8.33 (s, 1H), 7.99 (t, *J* = 5.6 Hz, 1H), 5.80 (d, *J* = 1.5 Hz, 1H), 5.55 (d, *J* = 6.3 Hz, 1H), 5.33 (br s, 1H), 4.04-4.28 (m, 3H), 3.96 (d, *J* = 8.1 Hz, 1H), 3.59-3.92 (m, 3H), 3.09-3.22 (m, 2H), 2.80-2.93 (m, 1H), 1.90 (s, 3H), 1.01-1.12 (m, 9H); $^{13}$C NMR (75 MHz, DMSO-$d_6$): δ 178.7, 168.7, 162.6, 153.5, 142.5, 107.0, 88.4, 83.8, 83.2, 69.3, 67.1, 58.9, 45.6, 33.0, 19.2, 19.1, 14.7, 8.6; LRMS (ESI TOF): *m/z* calcd for $C_{18}H_{29}N_4O_7$: 412.2, found 413.2 $[M + H]^+$.

**Compound S30**. To a pyridine solution (25 mL, anhydrous) of compound **S29** (1.1 g, 2.7 mmol), DMTCl (1.8 g, 5.4 mmol) was added. The reaction mixture was stirred at room temperature for 12 h. The reaction was quenched by adding methanol (0.3 mL) and stirring for 30 min at room temperature. Diluted the reaction mixture with ethyl acetate (50 mL) washed with saturated aqueous sodium bicarbonate (100 mL), brine (100 mL), dried ($Na_2SO_4$) and concentrated under reduced pressure. The crude product was purified by silica gel column chromatography (20 g) and eluted first with 0-50% ethyl acetate in the hexanes (5 column volume) and then 0-100% ethyl acetate in the dichloromethane to yield **S30** (1.7 g, 86%). $^{1}$H NMR (300 MHz, DMSO-d$_6$): δ 10.03 (br s, 1H), 7.98 (t, *J* = 5.7 Hz, 1H), 7.88 (br s, 1H), 7.40-7.47 (m, 2H), 7.23-7.37 (m, 7H), 6.91 (d, *J* = 9.0 Hz, 4H), 5.82 (d, *J* = 0.9 Hz, 1H), 5.62 (d, *J* = 7.9 Hz, 1H), 3.97-4.41 (m, 5H), 3.74 (s, 6H), 3.27-3.42 (m, 2H), 3.07-3.23 (m, 2H), 2.88 (br s, 1H), 1.50 (s, 3H), 1.00-1.21 (m, 9H); $^{13}$C NMR (75 MHz, DMSO-d$_6$) d: 175.8, 175.6, 168.7, 158.2, 144.5, 135.4, 135.1, 129.8, 129.7, 128.0, 127.9, 127.7, 126.8, 113.3, 89.0, 85.8, 82.7, 81.8, 69.4, 68.0, 59.7, 55.0, 33.0, 29.6, 19.2, 19.1, 14.7; LRMS (ESI TOF): *m/z* calcd for $C_{39}H_{47}N_4O_9$: 714.3, found 715.3 $[M + H]^+$.

**5'-*O*-DMT-2'-*O*-(NEA)-*N*-4-isobutyryl-5-methyl-cytidine-3'-phosphoramidite S31.** Compound **S30** (1.7 g, 2.3 mmol) and diisopropyl ammonium tetrazolide (0.4 g, 2.2 mmol) were placed in a flask with a stir bar and under reduced pressure over $P_2O_5$ for three days. The mixtures were suspended in anhydrous acetonitrile (15 mL). To this suspension, 2-cyanoethyl *N*,*N*,*N*′,*N*′-tetraisopropylphosphorodiamidite (1.2 mL, 3.5 mmol) was added. The reaction mixture was stirred at room temperature for 5 h and filtered. The filtrate was concentrated under reduced pressure. The residue was purified by silica gel column (primed with 1% V/V triethylamine in dichloromethane) and eluted with 50-70% ethyl acetate in a dichloromethane containing triethylamine (1% V/V) to yield **S31** (2.1 g, <95%). $^{31}$P NMR (121 MHz, DMSO-d$_6$) d: 150.18, 148.30; HRMS (ESI TOF): *m/z* calcd for $C_{48}H_{62}N_6O_{10}P$: 913.4343, found 913.4262 $[M - H]^-$.

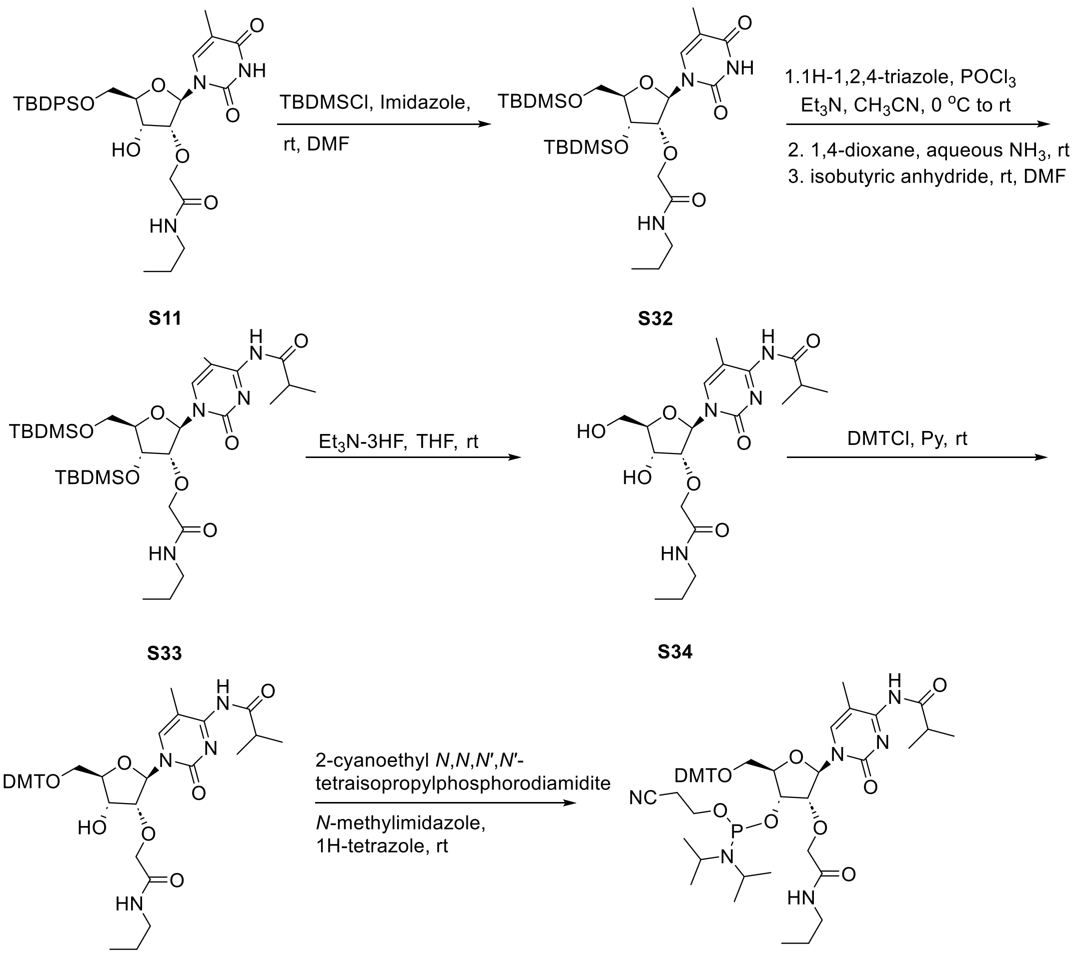

**Scheme S8**. Synthesis of 5'-*O*-DMT-2'-*O*-(NPA)-*N*-4-isobutyryl-5-methyl-cytidine-3'-phosphoramidite **S36**

**Compound S32**. To a solution of compound **S11** (6.8 g, 11.4 mmol) in DMF (76 mL) imidazole (4.6 g, 68.2mmol), *tert*-butyl-chlorodimethylsilane (6.9 g, 45.5 mmol) was added. The reaction mixture was stirred at room temperature for 12 h. MeOH (2 mL) was added and stirring continued for 30 min. After that the reaction mixture was diluted with ethyl acetate (100 mL) and washed with brine, dried ($Na_2SO_4$) and concentrated under reduced pressure. The residue was purified by silica gel column chromatography (0-5% methanol in a dichloromethane) to yield **S32** (5.8 g, 72%). $^1$H NMR (300 MHz, DMSO-$d_6$): δ 11.42 (s, 1H), 7.64 (dt, $J$ = 1.6, 8.0 Hz, 4H), 7.26-7.52

(m, 8H), 5.91 (d, $J$ = 4.7 Hz, 1H), 4.38 (t, $J$ = 4.9 Hz, 1H), 4.18 (t, $J$ = 5.0 Hz, 1H), 3.76-4.03 (m, 5H), 2.96-3.10 (m, 2H), 1.49 (s, 3H), 1.30-1.45 (m, 2H), 1.02 (s, 9H), 0.75-0.91 (m, 12H), 0.07 (s, 3H), 0.05 (s, 3H); $^{13}$C NMR (75 MHz, DMSO-$d_6$): δ 168.1, 163.7, 150.5, 135.8, 135.0, 132.6, 132.3, 128.0, 109.5, 86.8, 83.9, 80.3, 69.7, 69.0, 63.1, 54.9, 26.7, 25.8, 25.6, 22.3, 18.8, 17.7, 11.8, 11.2, -4.8, -5.1; LRMS (ESI TOF): *m/z* calcd for $C_{37}H_{56}N_3O_7Si_2$: 710.4, found 710.4 $[M + H]^+$.

**Compound S33**. $POCl_3$ (6.0 mL, 65.2 mmol) was slowly added to a suspension of 1,2,4-triazole (18.0 g, 261.0 mmol) in acetonitrile (160 mL) in an ice bath. The mixture was stirred at 0 $^o$C for 15 min and triethylamine (45.5 mL, 326.0 mmol) was added. The reaction mixture was stirred at 0 $^o$C (ice bath) for 30 min. A solution of compound **S32** (5.8 g, 8.2 mmol) in acetonitrile (36 mL) was added. The reaction mixture was removed from the ice bath and stirred at room temperature for 18 h. The reaction mixture was diluted with ethyl acetate (150 mL) and washed with water (100 mL), dried (Na2SO4, filtered and concentrated under reduced pressure to yield a triazolide analog of compound **S32** (6.2 g). LRMS (ESI TOF): *m/z* calcd for $C_{39}H_{56}N_6NaO_6Si_2$: 783.3, found 783.3 $[M + Na]^+$.  To a 1,4-dioxane (56 mL) solution of triazolide analog of compound **S32** (6.2 g, 8.2 mmol), aqueous ammonia (33.3 mL, 28-30 wt%) was added. The round bottom flask containing the reaction mixture was sealed with a septum and stirred at room temperature for eight h. The reaction mixture was concentrated under reduced pressure to yield 5-methylcytidine analog of compound **S32** (5.8 g). LRMS (ESI TOF): *m/z* calcd for $C_{37}H_{57}N_4O_6Si_2$: 709.3, found 709.3 $[M + 1]^+$. 5-Methylcytidine analog of compound **S32** (5.8 g, 8.2 mmol) from above was dried over $P_2O_5$ under reduced pressure overnight and dissolved in anhydrous DMF solution (81 mL, anhydrous). To this isobutyric anhydride (3.0 mL, 20.4 mmol) was added and stirred at room temperature for 5 h. The reaction was quenched by adding MeOH (2 mL) and stirring for 1o min. Diluted the reaction mixture with ethyl acetate (100 mL) and washed with brine (150 mL), dried

($Na_2SO_4$), filtered and concentrated under reduced pressure. The residue was purified by silica gel column chromatography (20-70% ethyl acetate in dichloromethane) to give **S33** (3.4 g, 54%). $^1$H NMR (300 MHz, DMSO-$d_6$): δ 10.19 (br s, 1H), 7.78 (s, 1H), 7.66 (ddd, *J* = 1.6, 7.8, 11.6 Hz, 4H), 7.54 (t, *J* = 5.9 Hz, 1H), 7.29-7.51 (m, 6H), 5.92 (d, *J* = 2.8 Hz, 1H), 4.35 (dd, *J* = 5.2, 6.7 Hz, 1H), 3.81-4.23 (m, 6H), 2.95-3.18 (m, 2H), 2.45-2.55 (m, 1H), 1.55 (s, 3H), 1.30-1.49 (m, 2H), 0.99-1.11 (m, 15H), 0.73-0.88 (m, 12H), 0.06 (s, 3H), 0.03 (s, 3H); $^{13}$C NMR (75 MHz, DMSO-d6): δ 168.3, 135.0, 134.9, 132.7, 132.4, 130.1, 130.0, 128.0, 127.9, 88.9, 83.4, 81.5, 69.4, 69.2, 62.6, 26.8, 25.5, 22.4, 19.1, 19.1, 19.0, 17.6, 13.6, 11.3, -4.7, -5.2; LRMS (ESI TOF): *m/z* calcd for $C_{41}H_{63}N_4O_7Si_2$: 779.4, found 779.4 $[M + H]^+$.

**Compound S34**. To a THF solution (40 mL) of compound **S33** (3.6 g, 4.6 mmol) triethylamine (2.2 mL, 15.9 mmol)) and triethylamine trihydrofluoride (2.3 mL, 13.8 mmol) were added. The mixture was stirred at room temperature for 12 h and the reaction mixture was concentratedunder reduced pressure. The residue thus obtained was purified by silica gel column chromatograph (0-20% MeOH in the dichloromethane) to yield **S34** (2.0 g, <95%). $^1$H NMR (300 MHz, DMSO-$d_6$): δ 10.19 (br s, 1H), 8.31 (s, 1H), 7.97 (t, *J* = 5.7 Hz, 1H), 5.79 (d, *J* = 1.7 Hz, 1H), 5.56 (d, *J* = 7.2 Hz, 1H), 5.33 (br s, 1H), 3.60-4.27 (m, 7H), 3.03-3.14 (m, 2H), 2.80-2.93 (m, 1H), 1.89 (s, 3H), 1.30-1.59 (m, 2H), 1.09 (s, 3H), 1.07 (s, 3H), 0.84 (t, *J* = 7.4 Hz, 3H); 13C NMR (75 MHz, DMSO-$d_6$): δ 168.9, 134.0, 130.6, 128.2, 107.0, 88.4, 83.8, 83.3, 69.3, 67.1, 58.9, 54.9, 45.6, 25.7, 22.4, 19.2, 19.2, 11.4, 8.8; LRMS (ESI TOF): *m/z* calcd for $C_{19}H_{31}N_4O_7$: 427.2, found 427.2 $[M + 1]^+$.

**Compound S35**. DMTCl (3.1 g, 9.2 mmol) was added to a solution of **S39** (1.97 g, 4.6 mmol), in anhydrous pyridine (45 mL) at 0 $^o$C in an ice bath. The reaction mixture was removed from the ice bath and stirred at room temperature for 12 h. The reaction was quenched by adding methanol (0.5 mL) and stirring at room temperature for 30 min. Subsequently, the reaction mixture was diluted with ethyl acetate (100 mL) washed with saturated aqueous sodium bicarbonate (100 mL), brine (100 mL), dried ($Na_2SO_4$), filtered, and concentrated under reduced pressure. The residue was purified by silica gel column chromatography (0-100% ethyl acetate in the dichloromethane) to give **S35** (2.7 g, 80%). $^1$H NMR (300 MHz, DMSO-d$_6$): δ 10.02 (br s, 1H), 7.97 (t, *J* = 5.8 Hz, 1H), 7.88 (br s, 1H), 7.38-7.51 (m, 2H), 7.14-7.38 (m, 7H), 6.91 (d, *J* = 9.0 Hz, 4H), 5.81 (s, 1H), 5.64 (d, *J* = 7.8 Hz, 1H), 3.94-4.39 (m, 5H), 3.74 (s, 6H), 3.26-3.43 (m, 2H), 3.08 (q, *J* = 6.5 Hz, 2H), 2.87 (br s, 1H), 1.50 (s, 3H), 1.36-1.48 (m, 2H), 1.09 (s, 3H), 1.07 (s, 3H), 0.84 (t, *J* = 7.4 Hz, 3H); $^{13}$C NMR (75 MHz, DMSO-d$_6$): δ 170.7, 169.2, 158.4, 144.7, 135.6, 135.3, 130.0, 129.9, 128.2, 127.9, 127.1, 113.5, 107.1, 89.4, 86.0, 83.0, 82.0, 78.9, 69.5, 69.4, 68.0, 61.9, 60.0, 55.2, 34.5, 33.9, 33.6, 29.6, 27.8, 23.1, 22.5, 20.9, 19.4, 19.2, 14.3, 11.5; LRMS (ESI TOF): *m/z* calcd for $C_{40}H_{49}N_4O_9$: 729.3, found 729.3 $[M + H]^+$.

**5’-*O*-DMT-2’-*O*-(NPA)-*N*-4-isobutyryl-5-methyl-cytidine-3’-phosphoramidite S36.** Compound **S35** (2.7 g, 3.7 mmol) and diisopropyl ammonium tetrazolide (0.6 g, 3.5 mmol) were placed in a flask with a stir bar and under reduced pressure over $P_2O_5$ overnight. The mixtures were suspended in anhydrous acetonitrile (24 mL) and 2-Cyanoethyl *N*,*N*,*N*′,*N*′-tetraisopropylphosphorodiamidite (1.8 mL, 5.5 mmol) was added. The reaction mixture was stirred at room temperature for 5 h and filtered. The filtrate was concentrated under reduced pressure. The residue was purified by silica gel column chromatography (primed with 1% triethylamine in dichloromethane) and eluted with 50-70% ethyl acetate in a dichloromethane containing triethylamine (1% V/V) to yield **S36** (3.4 g, <95%) was obtained as the desired product. $^{31}$P NMR (121 MHz, DMSO-d$_6$): δ 150.16, 148.32; HRMS (ESI TOF): *m/z* calcd for $C_{49}H_{64}N_6O_{10}P$: 927.4500, found 927.4420 $[M - H]^-$.

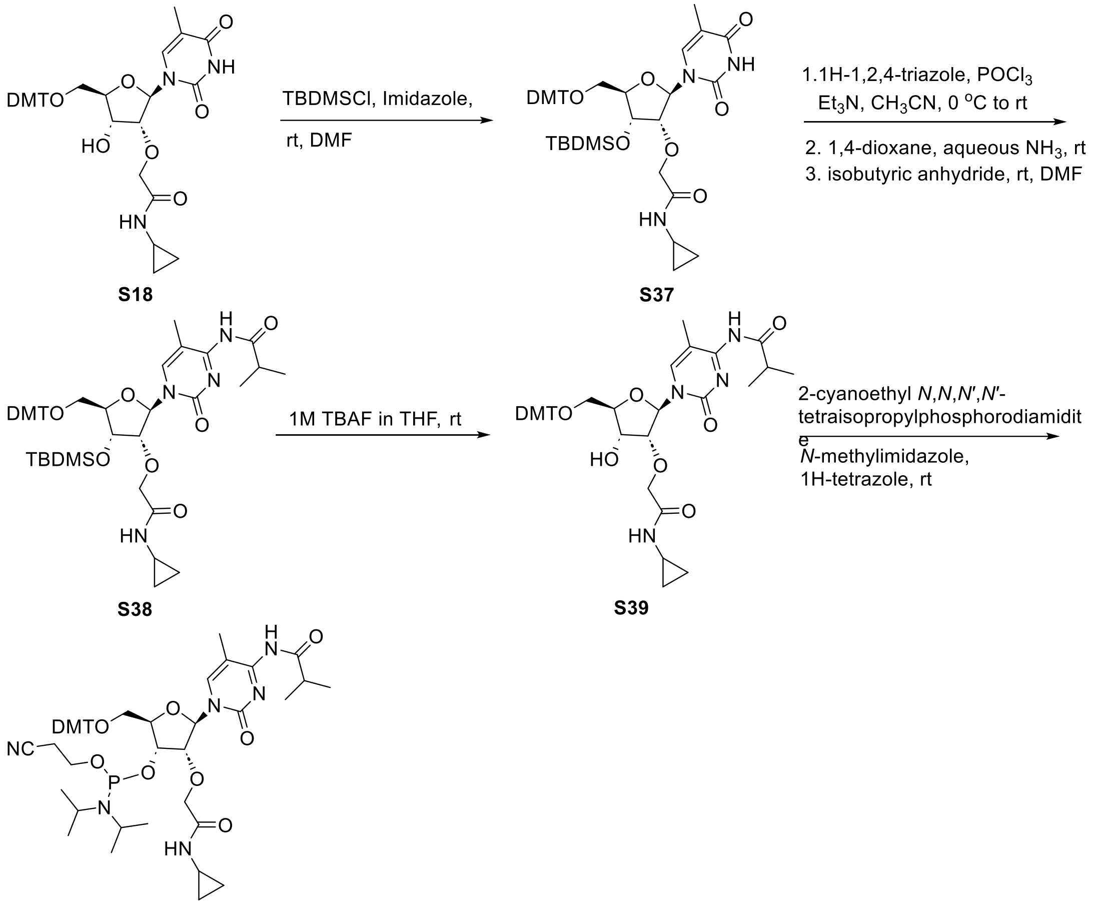

**Scheme S9**. Synthesis of 5'-*O*-DMT-2'-*O*-(NcPA)-*N*-4-isobutyryl-5-methyl-cytidine-3'-phosphoramidite **S40**

**Compound S37.** Compound **S18** (3.0 g, 4.6 mmol) was dissolved in anhydrous DMF (30 mL). To this, imidazole (1.9 g, 27.4 mmol) and tert-butyl-chlorodimethylsilane (2.8 g, 18.2 mmol) were added and stirred at room temperature for 12 h. The reaction mixture was diluted ethyl acetate and washed with saturated aqueous sodium bicarbonate (150 mL), brine (150 mL), dried ($Na_2SO_4$), filtered, and concentrated under reduced pressure. The residue was purified with a silica gel column chromatography (0-5% methanol in a dichloromethane) to yield **S37** (3.2 g, 91%). $^1$H NMR (300 MHz, DMSO-$d_6$): δ 11.42 (s, 1H), 7.48-7.60 (m, 2H), 7.20-7.43 (m, 9H), 6.89 (dd, $J$ = 1.3, 8.9 Hz, 4H), 5.86 (d, $J$ = 3.7 Hz, 1H), 4.38 (t, $J$ = 5.6 Hz, 1H), 4.14-4.21 (m, 1H), 3.91-4.09

(m, 3H), 3.73 (s, 6H), 3.15-3.37 (m, 2H), 2.62 (qt, $J$ = 3.9, 7.36 Hz, 1H), 1.43 (s, 3H), 0.75 (s, 9H), 0.53-0.66 (m, 2H), 0.34-0.44 (m, 2H), 0.03 (s, 3H), -0.06 (s, 3H); $^{13}$C NMR (75 MHz, DMSO-$d_6$): δ 169.6, 163.8, 158.2, 150.4, 144.4, 135.8, 135.2, 135.1, 129.8, 129.7, 127.9, 127.7, 126.9, 113.2, 109.3, 87.5, 86.0, 82.4, 80.8, 70.0, 69.2, 62.4, 55.1, 25.5, 22.0, 17.6, 11.8, 5.7, 5.6, -4.7, -5.3; LRMS (ESI, TOF): *m/z* calcd for $C_{42}H_{52}N_3O_9Si$: 770.3, found 770.3 $[M - H]^-$.

**Compound S38**. To an ice-cold suspension of 1,2,4-triazole (9.2 g, 133.0 mmol) in acetonitrile (70 mL) $POCl_3$ (3.1 mL, 33.4 mmol) was slowly. After stirring at 0 $^o$C (ice bath) for 15 min triethylamine (23.3 mL, 167.0 mmol) was added and stirring continued for an additional 30 min at 0 $^o$C. A solution of compound **S37** (3.2 g, 4.2 mmol) in anhydrous acetonitrile (20 mL) was added, and the reaction mixture was removed from the ice bath and stirring continued at room temperature for an additional 18 h. The reaction was quenched with water (20 mL) and diluted with ethyl acetate (200 mL) and washed with brine (200 mL), dried ($Na_2SO_4$), filtered, and concentrated under reduced pressure to yield a triazolide analog of compound **S38**. To a 1,4-dioxane (31 mL) solution of a triazolide analog of compound **S37** (3.4 g, 4.2 mmol) from above, aqueous ammonia (17.0 mL, 28-30 wt%) was added, sealed the reaction vessel, and stirred at room temperature for 3 h. The reaction mixture was concentrated under reduced pressure to yield a 5-methylcytidine analog of compound **S37** (3.2 g). The 5-methylcytidine analog of compound **S37** (3.2 g, 4.2 mmol) was dried over $P_2O_5$ under reduced pressure and then dissolved in anhydrous DMF (42 mL). Isobutyric anhydride (1.2 mL, 8.3 mmol) was added to this solution and stirred at room temperature for 20 h. The reaction was quenched by adding MeOH (1 mL) and stirring at room temperature for 10 min. Ethyl acetate (100 mL) was added to the reaction mixture, and the resulting solution was washed with brine (150 mL), dried ($Na_2SO_4$), filtered, and concentrated under reduced pressure. The residue was purified by silica gel column chromatography (0-80% ethyl acetate in dichloromethane) to give **S38** (2.1 g, 61%). $^{1}$H NMR (300 MHz, DMSO-$d_6$): δ 7.60 (s, 1H), 7.37-7.51 (m, 2H), 7.13-7.37 (m, 7H), 6.89 (dd, *J* = 1.8, 9.0 Hz, 4H), 5.84 (dd, *J* =1.7, 8.0 Hz, 1H), 4.28-4.44 (m, 1H), 3.93-4.23 (m, 4H), 3.73 (s, 6H), 3.25-3.54 (m, 3H), 3.15 (dt, *J* = 3.8, 10.78 Hz, 1H), 2.59-2.77 (m, 1H), 2.32-2.48 (m, 1H), 1.41-1.51 (m, 3H), 1.06 (s, 3H), 1.04 (s, 3H), 0.71 (s, 9H), 0.56-0.67 (m, 2H), 0.45-0.54 (m, 2H), 0.00 (s, 3H), -0.13 (s, 3H); $^{13}$C NMR (75 MHz, DMSO-$d_6$): δ 177.8, 169.8, 169.7, 165.6, 158.2, 155.0, 144.3, 144.3, 137.4, 135.2, 135.1, 129.8, 129.8, 127.9, 126.9, 113.2, 101.3, 88.7, 85.9, 82.0, 81.5, 69.8, 69.5, 68.5, 61.9, 55.8, 55.1, 33.1, 29.6, 25.5, 22.1, 18.9, 17.5, 13.0, 5.7, 5.4, -4.6, -5.4, ; LRMS (ESI TOF): *m/z* calcd for $C_{46}H_{61}N_4O_9Si$: 841.4, found 841.4 [M + H]+.

**Compound S39**. To a solution of compound **S38** (2.1 g, 2.5 mmol) in THF (25 mL), tetrabutylammonium fluoride in THF (3.82 mL, 1 M) was added. The reaction mixture was stirred at room temperature for 16 h and concentrated under reduced pressure. The residue was dissolved in ethyl acetate (100 mL), washed with brine (100 mL), dried ($Na_2SO_4$), filtered, and concentrated under reduced pressure. The residue was purified by silica gel column chromatography (0-50% acetone in dichloromethane) to yield **S39** (1.5 g, 78%). $^{1}$H NMR (300 MHz, DMSO-$d_6$): δ 7.91-8.06 (m, 1H), 7.87 (s, 1H), 7.39-7.46 (m, 2H), 7.20-7.37 (m, 8H), 6.91 (d, *J* = 9.0 Hz, 4H), 5.79 (d, *J* = 0.9 Hz, 1H), 5.66 (d, *J* = 7.8 Hz, 1H), 4.06-4.37 (m, 4H), 3.90-4.02 (m, 1H), 3.74 (s, 6H), 3.16-3.43 (m, 2H), 2.77-3.01 (m, 1H), 2.68 (qt, *J* = 3.9, 7.4 Hz, 1H), 1.49 (s, 3H), 1.09 (s, 3H), 1.07 (s, 3H), 0.59-0.69 (m, 2H), 0.41-0.53 (m, 2H); $^{13}$C NMR (75 MHz, DMSO-$d_6$): δ 170.2, 158.2, 144.5, 135.4, 135.1, 129.8, 129.7, 128.0, 127.7, 126.8, 113.3, 85.8, 82.8, 81.8, 69.3, 67.9, 61.7, 55.0, 53.2, 32.1, 30.7, 29.6, 22.0, 20.0, 19.2, 19.1, 13.9, 13.7, 5.6, 5.5; LRMS (ESI TOF): *m/z* calcd for $C_{40}H_{47}N_4O_9$: 727.3, found 727.3 $[M + H]^+$.

**5’-*O*-DMT-2’-*O*-(NcPA)-*N*-4-isobutyryl-5-methyl-cytidine-3’-phosphoramidite S40.** Compound **S39** (1.5 g, 2.0 mmol) and diisopropyl ammonium tetrazolide (0.3 g, 1.9 mmol) were placed in a flask with a stir bar dried over $P_2O_5$ under reduced pressure for 15 h. The mixture was suspended in anhydrous acetonitrile (13 mL) and 2-cyanoethyl *N*,*N*,*N′*,*N′*-tetraisopropylphosphorodiamidite (1.0 mL, 3.0 mmol) was added. The reaction mixture was stirred at room temperature for 4 h and filtered. The filtrate was concentrated under reduced pressure and the residue product was purified by silica gel column (eluted with 50-80% ethyl acetate in hexanes containing 1% V/V triethylamine) to yield **S40** (1.8 g, <95%). $^{31}$P NMR (121 MHz, DMSO-d$_6$): δ 150.14, 148.39; HRMS (ESI TOF): *m/z* calcd for $C_{49}H_{62}N_6O_{10}P$: 925.4343, found 925.4264 [M - H]$^-$.

**Compound S41**. Compound **S21** (3.0 g, 4.5 mmol) was dissolved in anhydrous DMF (30 mL). Imidazole (1.8 g, 26.8 mmol) and *tert*-butyl-chlorodimethylsilane (2.7 g, 17.9 mmol) were added. The reaction mixture was stirred at room temperature for 12 h. Th reaction was quenched by adding MeOH (2 mL) and stirring for 30 min. Ethyl acetate (100 mL) was added, and the organic layer was washed with brine (100 mL), dried ($Na_2SO_4$), filtered, and concentrated under reduced pressure. The residue was purified by silica gel column chromatography (0-5% methanol in dichloromethane) to yield **S41** (3.4 g, 97%). $^1$H NMR (300 MHz, DMSO-d$_6$): δ 11.43 (s, 1H), 7.59 (s, 1H), 7.54 (t, *J* = 5.8 Hz, 1H), 7.37-7.44 (m, 2H), 7.21-7.35 (m, 7H), 6.89 (dd, *J* = 1.5, 9.0 Hz, 4H), 5.87 (d, *J* = 3.33 Hz, 1H), 4.34-4.45 (m, 1H), 3.92-4.19 (m, 4H), 3.68-3.83 (m, 6H), 3.29-3.40 (m, 2H), 2.90-3.07 (m, 2H), 1.44 (d, *J* = 0.90 Hz, 3H), 0.85-0.95 (m, 1H), 0.70-0.79 (m, 9H), 0.30-0.42 (m, 2H), 0.10-0.20 (m, 2H), 0.03 (s, 3H), -0.06 (s, 3H); $^{13}$C NMR (75 MHz, DMSO-d$_6$) δ 168.2, 163.8, 158.2, 150.4, 144.4, 135.8, 135.2, 135.1, 129.8, 129.7, 127.9, 127.7, 126.9, 113.2,

109.3, 87.7, 86.0, 82.3, 80.9, 69.9, 69.2, 62.4, 55.1, 42.5, 25.5, 17.6, 11.9, 10.8, 3.2, -4.7, -5.3; LRMS (ESI TOF): *m/z* calcd for $C_{43}H_{54}N_3O_9Si$: 784.4, found 784.3 $[M - H]^-$.

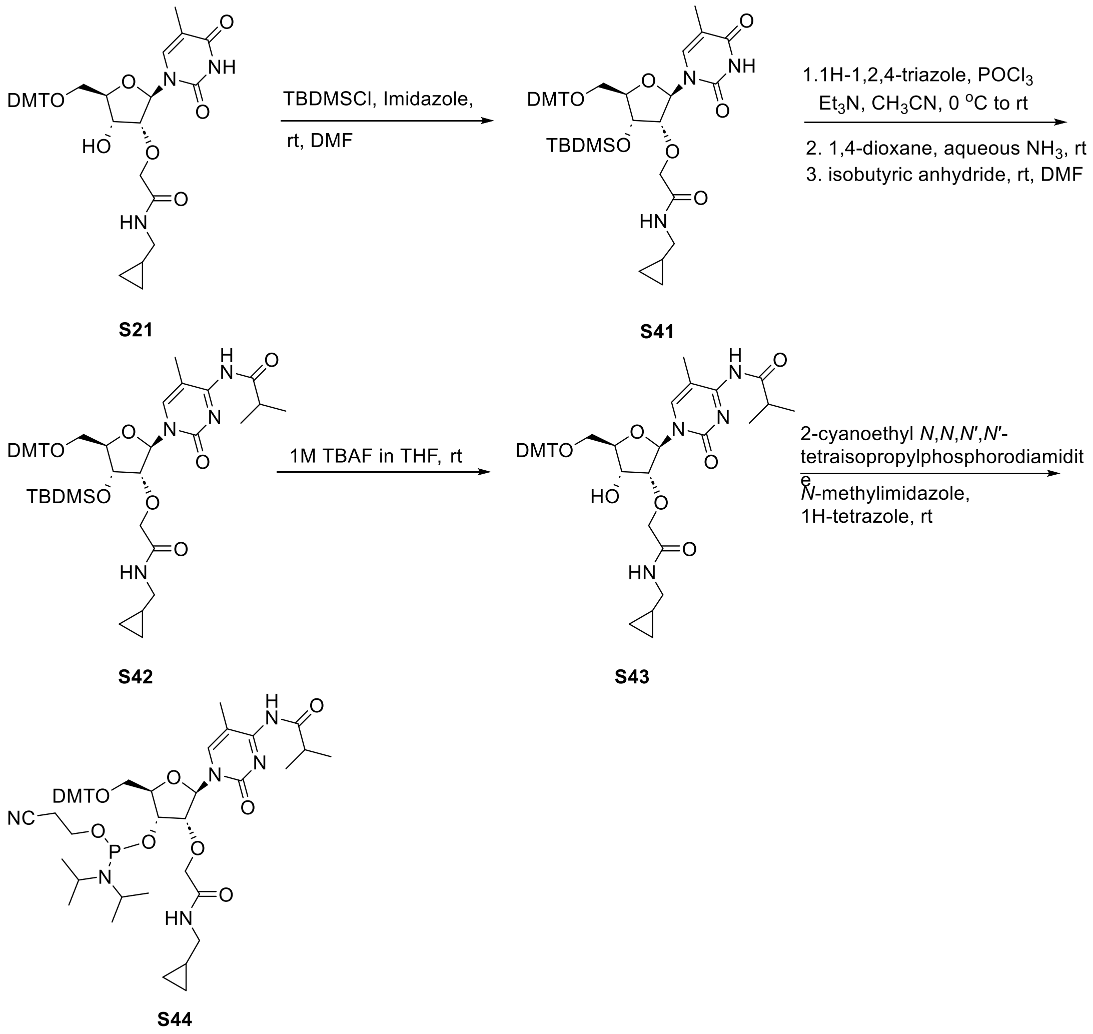

**Scheme S10**. Synthesis of 5'-*O*-DMT-2'-*O*-(McPA)-*N*-4-isobutyryl-5-methyl-cytidine-3'-phosphoramidite **S44**

**Compound S42.** To 1,2,4-triazole (9.6 g, 138.0 mmol) suspension in anhydrous acetonitrile (75 mL) $POCl_3$ (3.2 mL, 34.6 mmol) was slowly added at 0 °C (ice bath). After stirring for 15 min at triethylamine (24.1 mL, 173.0 mmol) was added and stirring continued for 30 min at 0 °C. A solution of compound **S41** (3.4 g, 4.3 mmol) in anhydrous acetonitrile (20 mL) was added. The reaction mixture was removed from the ice bath and stirred for five hours at room temperature. Ethyl acetate (100 mL) was added, and the organic phase was washed with brine (100 mL), dried ($Na_2SO_4$), filtered, and concentrated under reduced pressure to yield a triazolide analog of **S41**. The triazolide analog of **S41**(3.6 g, 4.3 mmol) from above was dissolved in 1,4-dioxane (33 mL), and aqueous ammonia (17.5 mL, 28-30 wt%) was added. The reaction vessel was sealed tightly and after stirring at room temperature for 2 h it was concentrated under reduced pressure to yield a 5-methylcytidine analog of compound **S41**. After drying the 5-methylcytidine analog of compound **S42** (3.4 g, 4.3 mmol), from above over $P_2O_5$ under reduced pressure overnight, it was dissolved in anhydrous DMF (44 mL), and isobutyric anhydride (1.3 mL, 8.6 mmol) was added. After stirring the reaction mixture for 20 h at room temperature, it was diluted with ethyl acetate (100 mL), and the organic phase was washed with brine (100 mL), dried ($Na_2SO_4$), filtered, and concentrated under reduced pressure. The residue was purified by silica gel column chromatography (0-30% acetone in dichloromethane) to yield **S42** (2.2 g, 60%). $^1$H NMR (300 MHz, DMSO-$d_6$): δ 8.02 (br s, 1H), 7.61 (t, *J* = 6.0 Hz, 1H), 7.37-7.48 (m, 2H), 7.14-7.37 (m, 8H), 6.90 (dd, *J* = 2.1, 9.0 Hz, 4H), 5.88 (d, *J* = 1.15 Hz, 1H), 4.38-4.46 (m, 1H), 4.02-4.32 (m, 4H), 3.69-3.82 (m, 6H), 3.48 (d, *J* = 9.5 Hz, 1H), 3.19 (dd, *J* = 3.8, 10.9 Hz, 1H), 2.80-3.14 (m, 3H), 1.50 (s, 3H), 1.09 (s, 3H), 1.07 (s, 3H), 0.89-1.02 (m, 1H), 0.58-0.77 (m, 9H), 0.29-0.45 (m, 2H), 0.10-0.24 (m, 2H), 0.01 (s, 3H), -0.11 (s, 3H); LRMS (ESI TOF): *m/z* calcd for $C_{47}H_{61}N_4O_9Si$: 853.5, found 853.5 $[M - H]^-$.

**Compound S43**. To a solution compound **S42** (2.2 g, 2.6 mmol) in THF (27 mL) tetrabutylammonium fluoride in THF (3.9 mL, 3.9 mmol) was added. The reaction mixture was stirred at room temperature for 16 h and concentrated under reduced pressure. The residue was dissolved in ethyl acetate (100 mL) and the organic phase was washed with brine (100 mL), dried ($Na_2SO_4$), filtered, and concentrated under reduced pressure. The residue was purified by silica gel column chromatography (0-100% ethyl acetate in a dichloromethane) to yield **S43** (1.7 g, 85%). $^1$H NMR (300 MHz, DMSO-d$_6$): δ 8.07 (t, *J* = 5.8 Hz, 1H), 7.88 (s, 1H), 7.18-7.50 (m, 10H), 6.91 (d, *J* = 9.0 Hz, 4H), 5.82 (d, *J* = 1.0 Hz, 1H), 5.64 (d, *J* = 7.6 Hz, 1H), 3.99-4.38 (m, 5H), 3.74 (s, 6H), 3.28-3.45 (m, 2H), 2.96-3.08 (m, 2H), 2.78-2.94 (m, 1H), 1.50 (s, 3H), 1.09 (s, 3H), 1.07 (s, 3H), 0.85-1.00 (m, 1H), 0.34-0.45 (m, 2H), 0.12-0.22 (m, 2H); $^{13}$C NMR (75 MHz, DMSO-d$_6$): δ 168.8, 158.2, 144.5, 135.4, 135.1, 129.8, 129.7, 128.0, 127.7, 126.8, 113.3, 89.0, 85.8, 82.8, 81.8, 69.4, 68.0, 61.8, 55.0, 42.5, 19.2, 19.1, 10.8, 3.2; LRMS (ESI TOF): *m/z* calcd for $C_{41}H_{49}N_4O_9$: 741.3, found 741.3 $[M + H]^+$.

**5'-*O*-DMT-2'-*O*-(McPA)-*N*-4-isobutyryl-5-methyl-cytidine-3'-phosphoramidite S44.** Compound **S43** (1.7 g, 2.2 mmol) and diisopropyl ammonium tetrazolide (0.4 g, 2.1 mmol) were placed in a flask with a stir bar and dried over $P_2O_5$ under reduced pressure for 15 h. The mixture was suspended in anhydrous acetonitrile (15 mL) and 2-cyanoethyl *N*,*N*,*N′*,*N′*-tetraisopropylphosphorodiamidite (1.1 mL, 3.3 mmol) was added. The reaction mixture was stirred at room temperature for 4 h and filtered. The filtrate was concentrated under reduced prssure. The residue was purified by a silica gel column chromatography and eluted with 50-100% ethyl acetate in hexanes containing triethylamine (1% V/V) to yield **S44** (1.8, 86%). $^{31}$P NMR (121 MHz, DMSO-d$_6$): δ 150.13, 148.40; HRMS (ESI TOF): *m/z* calcd for $C_{50}H_{64}N_6O_{10}P$: 939.4500, found 939.4422$[M - H]^-$.
